# An ALE meta-analytic review of musical expertise

**DOI:** 10.1101/2021.03.12.434473

**Authors:** Antonio Criscuolo, Victor Pando-Naude, Leonardo Bonetti, Peter Vuust, Elvira Brattico

## Abstract

Through long-term training, music experts acquire complex and specialized sensorimotor skills, which are paralleled by continuous neuro-anatomical and -functional adaptations. The underlying neuroplasticity mechanisms have been extensively explored in decades of research in music, cognitive, and translational neuroscience. However, the absence of a comprehensive review and quantitative meta-analysis prevents the plethora of variegated findings to ultimately converge into a unified picture of the neuroanatomy of musical expertise. Here, we performed a comprehensive neuroimaging meta-analysis of publications investigating neuro-anatomical and -functional differences between musicians (M) and non-musicians (NM). Eighty-four studies were included in the qualitative synthesis. From these, 58 publications were included in coordinate-based meta-analyses using the anatomic/activation likelihood estimation (ALE) method. This comprehensive approach delivers a coherent cortico-subcortical network encompassing sensorimotor and limbic regions bilaterally. Particularly, M exhibited higher volume/activity in auditory, sensorimotor, interoceptive, and limbic brain areas and lower volume/activity in parietal areas as opposed to NM. Notably, we reveal topographical (dis-)similarities between the identified functional and anatomical networks and characterize their link to various cognitive functions by means of meta-analytic connectivity modelling. Overall, we effectively synthesized decades of research in the field and provide a consistent and controversies-free picture of the neuroanatomy of musical expertise.

## 1. Introduction

Decades of research in psychology, cognitive and translational neuroscience have attempted to deepen our understanding of the cognitive and neural processes which allow individuals to reach high levels of mastery within a given domain. For instance, expert musicians (M) seem to develop, through long-term training, complex and specialized auditory and sensorimotor skills, which enable them to achieve the highest levels of performance in playing a musical instrument (thus, the label of “virtuoso”)^1–3^. Notably, such debate extends to the association between the acquisition of such fascinating skills and the continuous neuro-anatomical and -functional changes in auditory, motor and higher-order cognitive control regions since childhood^3–7^. Furthermore, other non-music-specific cognitive functions seem to benefit from such trainings, as revealed by increased performance in working memory, intelligence, executive functions and inhibitory control tests^5,8–10^. These effects may represent, on the one hand, ‘neuroplastic’ adaptations to environmental demands and successful gene-environment interactions^2,11^. On the other hand, many background and environmental factors may influence observed neurocognitive changes, as we shall discuss later.

Neuroplasticity generally refers to the brain’s capacity to modulate its anatomical and functional features during maturation, learning, skill acquisition, environmental challenges, or pathology. Interestingly, these effects seem to be salient enough to be also observed macroscopically with magnetic resonance imaging (MRI). However, whole-brain analyses in humans often fail to convey a link between complex behaviour and neuroplastic mechanisms. Such difficulty might emerge due to methodological and sample differences. For instance, background variables such as genome, psychological (motivation, reward), and socio-economical characteristics are not always considered^12,13^ but may determine the predisposition to engage in specialized trainings (e.g., music or dance) and/or to possess fine-grained sensory processing and sensorimotor skills which are independent of, or precede the training. Consequently, the adoption of cross-sectional designs, as opposed to longitudinal designs, may lead to spurious observations of training-related neuroplasticity mechanisms as they may fail to isolate background and confounding factors from the ‘true’ training-related effects. In contrast, longitudinal studies allow to zoom into individual differences and to follow specific trajectories of neurocognitive development. However, longitudinal research is rather sparse^3,6^.

A common approach adopted to strengthen the link between observed neuro-anatomical and - functional changes and specialized trainings in cross-sectional designs is to quantify their correlation. Thus, the relation with the length of specialized trainings, is interpreted as evidence for a link between expertise and neuroplasticity (in various populations from athletes, chess-players, golfers^14^ and musicians^5^). These correlational approaches are usually further extended to investigate the link between specific trainings and higher-order cognitive functions^5^. Thus, improved intelligence, working memory and executive functions are commonly associated with duration and intensity of music training^5,8–10^. These studies should be interpreted with caution as they not necessarily allow to dissociate pre-existing and training-independent neurocognitive differences from real neuroplasticity effects. Additionally, musical training represents a stimulating experience which engages highly-specialized perceptual, motor, emotional and higher-order cognitive abilities^1^, ranging from multimodal (auditory, visual, motor) sensory perception, integration, predictions and fine movement control^15,16^. Furthermore, it stimulates mnemonic processes associated with the acquisition of long and complex bimanual finger sequences, as well as fine-grained auditory perception (absolute pitch)^17^. Thus, the daily and intensive training of such complex, varied and specialized sensory, sensorimotor and higher-order cognitive skills represents a very appealing scenario to investigate neuroplasticity mechanisms and to monitor the continuous underlying neuro-anatomical and -functional adaptations^3,18^.

Anatomical and functional resonance imaging studies (fMRI) have correlated audio-motor, parietal and occipital brain structures (the dorsal stream) to the ability to play an instrument via automatic and accurate associations between motor sequences and auditory events leading to multimodal predictions^15^, the simultaneous integration of multimodal auditory, visual and motor information, and fine-grained skills in auditory perception, kinaesthetic control, visual perception, and pattern recognition^16^. Also, dorsolateral prefrontal structures, basal ganglia and mesial temporal structures have further been related to musicians’ ability to memorize long and complex bimanual finger sequences and to translate musical symbols into motor sequences^17^. Importantly, however, music experts do not show an overall pattern of increased structure and/or task-based functional activity. While surely appealing, such assumption would narrow down the complexity of the rather heterogenous neurocognitive mechanisms of musical expertise^19,20^. Hence, some works reported co-occurrent patterns of increase and decrease of GM volumes when comparing musicians to non-musicians^21,22^, and other research highlighted negative correlations between GM volumes and musical expertise^22,23^. Similarly, investigations focusing on WM reported increased fractional anisotropy (i.e., a measure of WM integrity) and diffusivity in expert musicians as compared to non-musicians in the corticospinal tract^24,25^, internal capsule bundles^26^ and corpus callosum^27^, while others found reduced fractional anisotropy and increased radial diffusivity^28^. fMRI studies are similarly characterized by variegated observations, with musicianship being exclusively associated with stronger activity in, e.g., either premotor cortices^29^, right auditory cortex^30^, or prefrontal cortex^31^, ultimately failing to converge into a common functional network for music expertise.

Despite a growing interest in the topic, we here highlight that there has never been a quantitative meta-analytic attempt to summarize existing findings and provide a unified picture of the neuroanatomy of musical expertise. To address this limitation in the field, we conducted a comprehensive coordinate-based meta-analysis (CBMA) of (f)MRI studies using the anatomic/activation likelihood estimation (ALE) method^32^, to investigate the neuro-anatomical and - functional signatures associated with musical training on healthy humans. Specifically, we first provide a detailed overview of the studies included and their methods, paradigms, sample details and backgrounds so to guide the reader into a critical consideration of the results. Then, we characterize the topographical (dis)similarities between the identified functional and anatomical networks and link them to various cognitive functions by means of meta-analytic connectivity modelling (MACM).

## 2. Results

A total of 1169 records was identified through database searching, and 679 records were initially screened by title and abstract after removing duplicates. Next, 145 articles were assessed for eligibility in the full-text screening stage. From these, 84 studies fulfilled criteria for eligibility and were thus included in the qualitative synthesis. Finally, from the 84 studies, only 58 reported results in stereotactic coordinates (foci), either Talairach or Montreal Neurological Institute (MNI) three-dimensional-coordinate system which were therefore included in the quantitative synthesis (ALE meta-analyses) (**Supplementary Figure 1**).

### 2.1. Characteristics of studies

Details of the studies included in our work are provided in **Table 1**. Eighty-four publications met inclusion criteria and were included in the qualitative synthesis which was comprised of 3005 participants, with 1581 musicians (M) and 1424 non-musicians (NM). Eighteen studies (21%) included amateur musicians, and only 7 studies (8.3%) reported absolute pitch possessors (n = 97). Musical instruments were reported in most of the studies (81%): piano or keyboard (62%), string instruments (41%), wind instruments (26%), percussion instruments (17%), voice (8%), and 19% studies failed to report musicians’ instrument. Years of education was described only in 8% of the included studies. Years of musical training was reported in 63% of the studies, with a mean of 15.6 ± 5.9 years. The age of onset of musical training was reported in 49% of the studies, with a mean of 7.4 ± 2.3 years old. Weekly hours of training were reported in 32% of the studies, with a mean of 16.7 ± 8.9 hours per week.

**Table 1.**
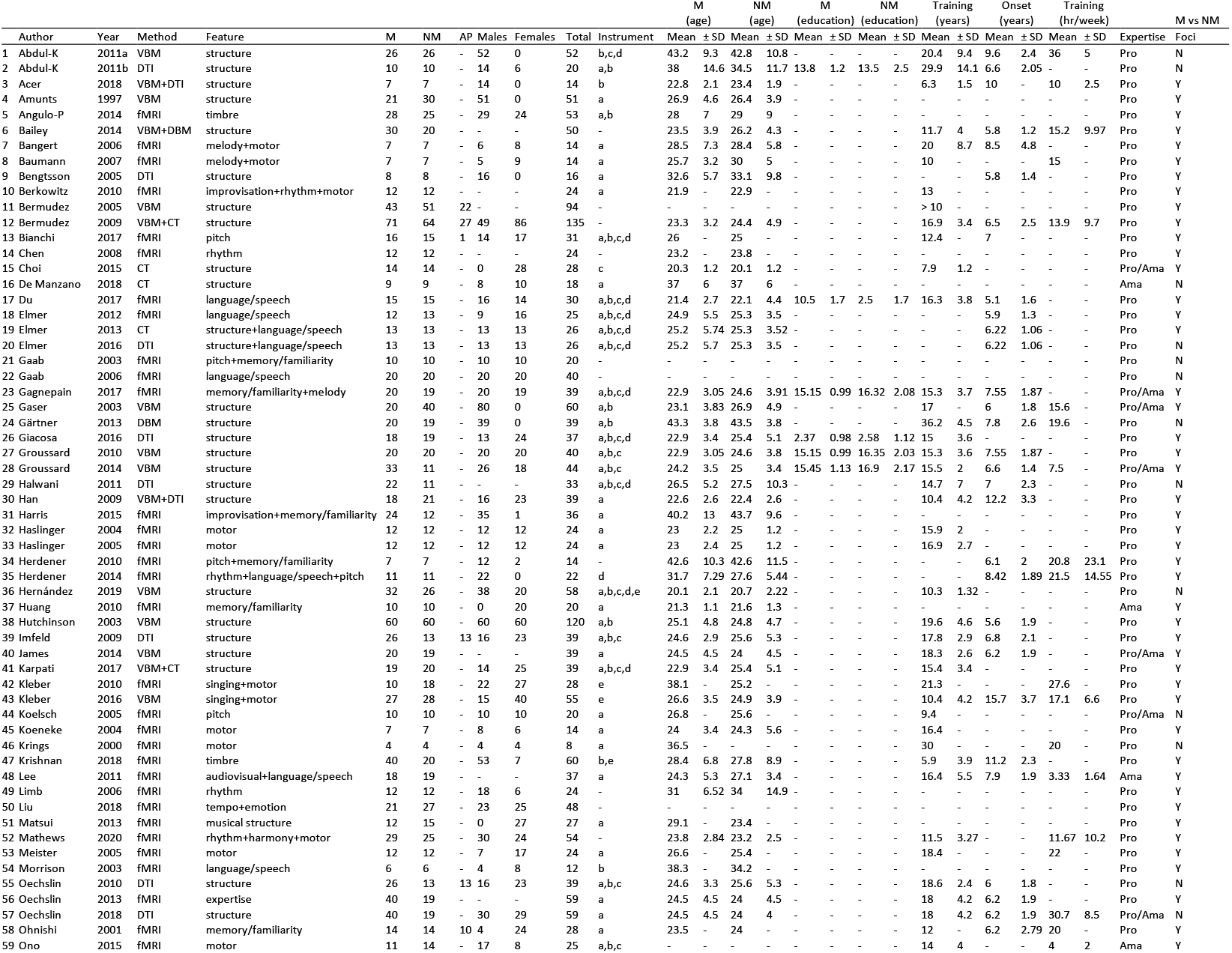
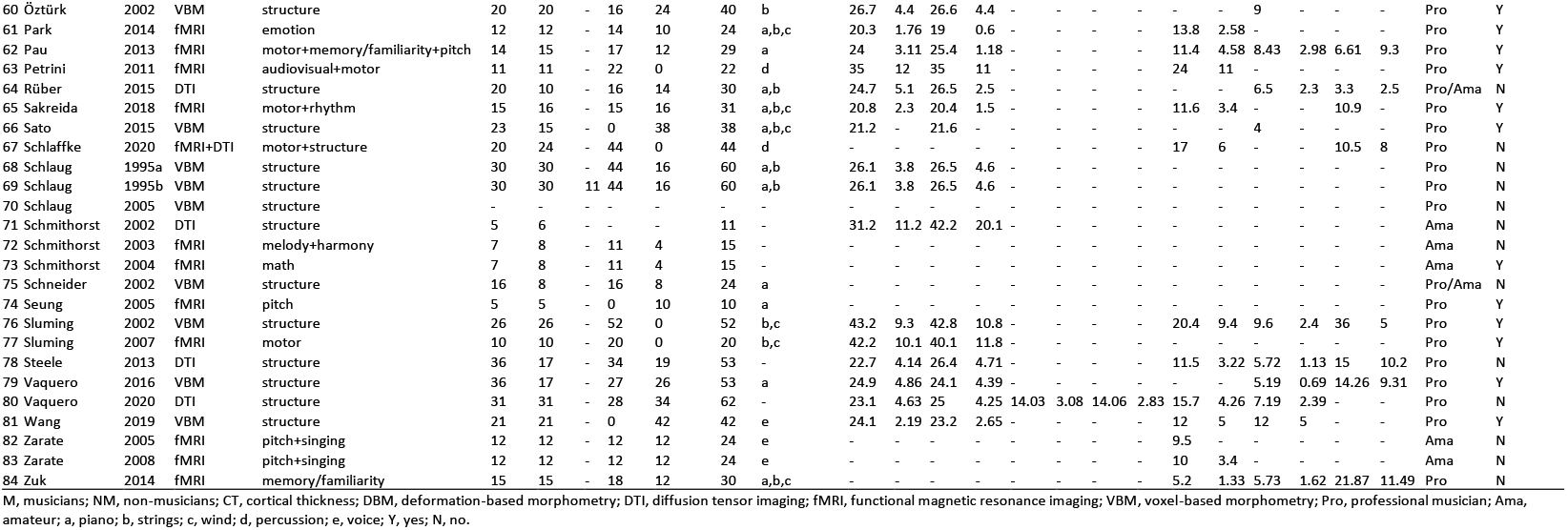
Characteristics of studies included in qualitative synthesis.

### 2.2. MRI quality

MRI quality of the included studies in the meta-analysis was assessed following a set of guidelines for the standardized reporting of MRI studies^33,34^. All studies included in the qualitative synthesis (n=84) reported their MRI design, software package and image acquisition, pre-processing, and analyses. Overall, all the studies followed good MRI practices. Neuroimaging data was acquired in either 1.5 T (39%), or 3 T (56%) scanners, while 5% of studies did not report the magnetic field strength. MRI scanners included Siemens (40%), General Electric (25%), Philips (21%), Bruker (5%), while 7% did not report it. Analysis methods included fMRI (52%), VBM (29%), DTI (18%), and CT (10%). Finally, 84% of studies included GM analyses, while 23% included WM analyses (**Supplementary Tables 1, 2, and 3**).

### 2.3. ALE meta-analysis

The quantitative synthesis of the primary outcome included 58 publications with 675 foci, 79 experiments and a total of 2,780 participants. Separate ALE meta-analyses were conducted in GingerALE^35^ for structural and functional foci, focusing on the comparison between M and NM.

#### 2.3.1. Structural studies

The structural ALE meta-analysis included 33 experiments and 1515 participants. The contrast M > NM in GM resulted in significant peak clusters in the bilateral superior temporal gyrus (primary auditory cortex), including the bilateral Heschl’s gyrus and planum temporale, and the postcentral gyrus (somatosensory cortex, SI), including area 4a of the primary motor cortex (M1 4a). Conversely, the comparison NM > M in GM resulted in a significant peak cluster located in the right precentral gyrus (primary motor cortex, M1). In WM, musicians showed larger tracts of the internal capsule bundle (extending to the thalamus) and corticospinal tract. No significant clusters were identified in the comparison NM>M for WM (**Figure 1**, **Table 2**).

**Figure 1.**
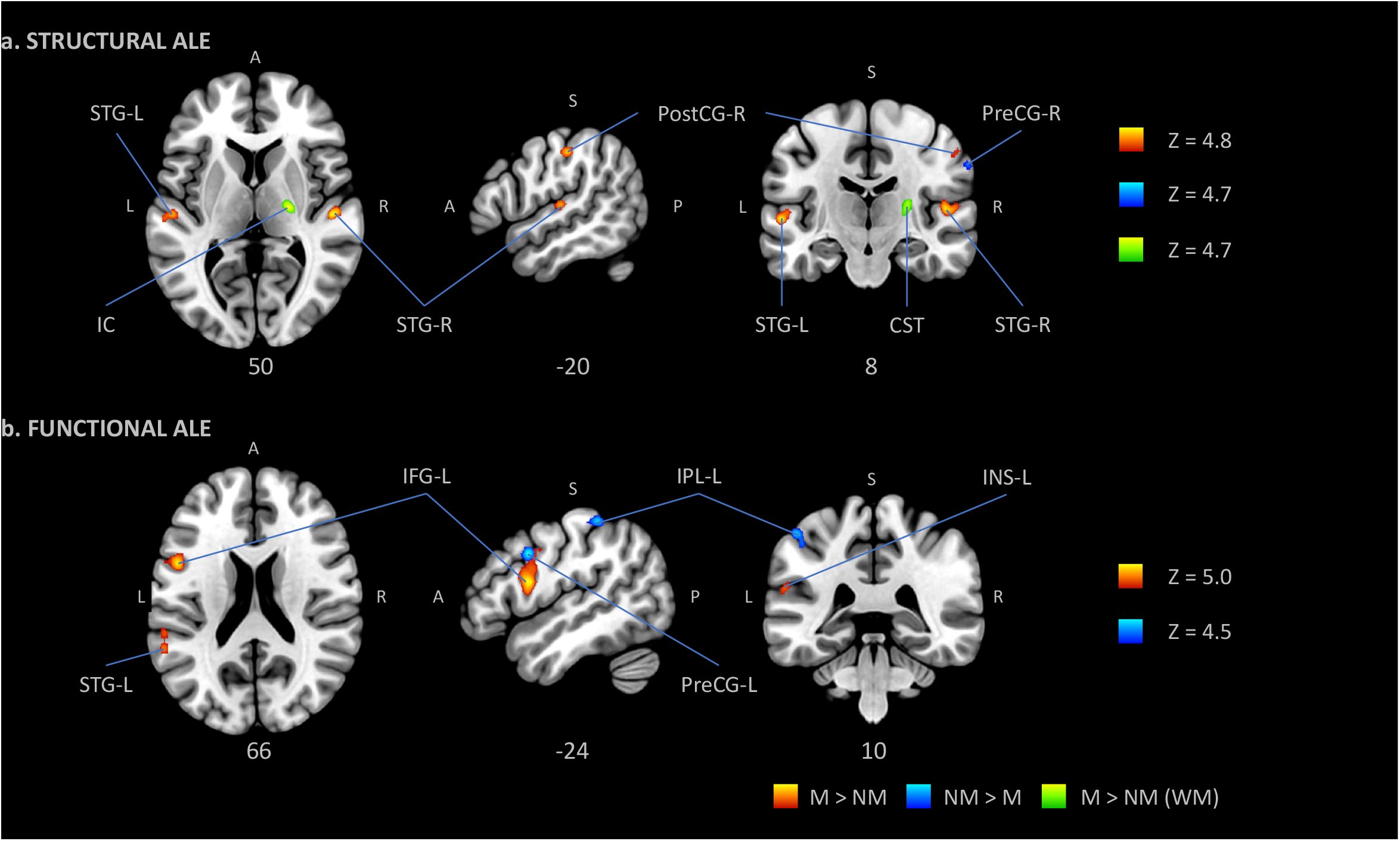
Anatomic likelihood estimation meta-analytic results for studies comparing brain structure and function between M and NM at cluster level inference p < 0.05 (FWE). The primary outcome included ALE meta-analysis of the contrast M vs NM for structural and functional modalities, independently. M > NM = higher volume/activity in musicians; NM > M = lower volume/activity in musicians; GM, grey matter; WM, white matter; L, left; R, right; A, anterior; P, posterior; Z, peak Z-value; IC, internal capsule; INS, insula; IPL, inferior parietal lobule; PostCG, postcentral gyrus (primary somatosensory cortex, or S1); PreCG, precentral gyrus (primary motor cortex, or M1); STG, superior temporal gyrus (primary auditory cortex).

**Table 2.**
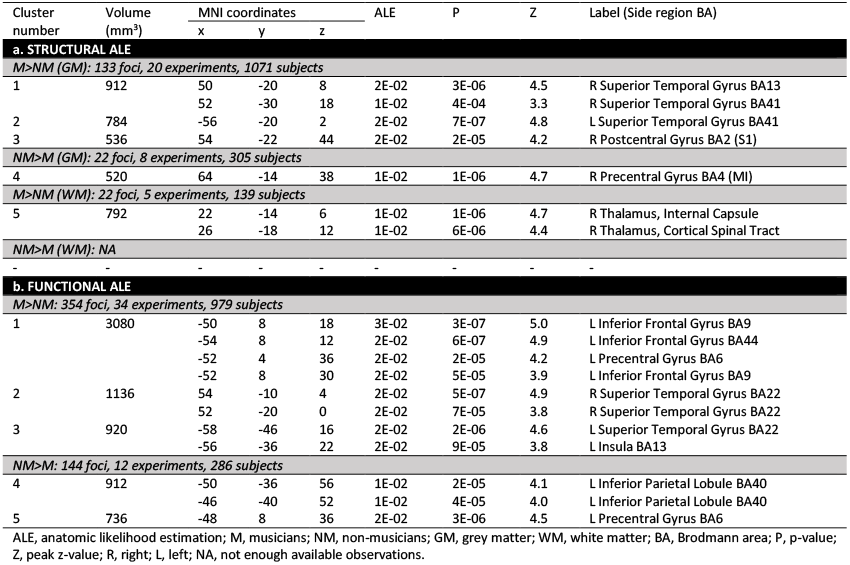
ALE meta-analytic results for structural and functional studies comparing M vs NM at cluster level inference p < 0.05 (FWE).

| Cluster number | Volume (mm³) | MNI coordinates |  |  | ALE | P | Z | Label (Side region BA) |
| --- | --- | --- | --- | --- | --- | --- | --- | --- |
|  |  | x | y | z |  |  |  |  |
| a. STRUCTURAL ALE |  |  |  |  |  |  |  |  |
| M>NM (GM): 133 foci, 20 experiments, 1071 subjects |  |  |  |  |  |  |  |  |
| 1 | 912 | 50 | -20 | 8 | 2E-02 | 3E-06 | 4.5 | R Superior Temporal Gyrus BA13 |
|  |  | 52 | -30 | 18 | 1E-02 | 4E-04 | 3.3 | R Superior Temporal Gyrus BA41 |
| 2 | 784 | -56 | -20 | 2 | 2E-02 | 7E-07 | 4.8 | L Superior Temporal Gyrus BA41 |
| 3 | 536 | 54 | -22 | 44 | 2E-02 | 2E-05 | 4.2 | R Postcentral Gyrus BA2 (S1) |
| NM>M (GM): 22 foci, 8 experiments, 305 subjects |  |  |  |  |  |  |  |  |
| 4 | 520 | 64 | -14 | 38 | 1E-02 | 1E-06 | 4.7 | R Precentral Gyrus BA4 (MI) |
| M>NM (WM): 22 foci, 5 experiments, 139 subjects |  |  |  |  |  |  |  |  |
| 5 | 792 | 22 | -14 | 6 | 1E-02 | 1E-06 | 4.7 | R Thalamus, Internal Capsule |
|  |  | 26 | -18 | 12 | 1E-02 | 6E-06 | 4.4 | R Thalamus, Cortical Spinal Tract |
| NM>M (WM): NA |  |  |  |  |  |  |  |  |
| - | - | - | - | - | - | - | - | - |
| b. FUNCTIONAL ALE |  |  |  |  |  |  |  |  |
| M>NM: 354 foci, 34 experiments, 979 subjects |  |  |  |  |  |  |  |  |
| 1 | 3080 | -50 | 8 | 18 | 3E-02 | 3E-07 | 5.0 | L Inferior Frontal Gyrus BA9 |
|  |  | -54 | 8 | 12 | 2E-02 | 6E-07 | 4.9 | L Inferior Frontal Gyrus BA44 |
|  |  | -52 | 4 | 36 | 2E-02 | 2E-05 | 4.2 | L Precentral Gyrus BA6 |
|  |  | -52 | 8 | 30 | 2E-02 | 5E-05 | 3.9 | L Inferior Frontal Gyrus BA9 |
| 2 | 1136 | 54 | -10 | 4 | 2E-02 | 5E-07 | 4.9 | R Superior Temporal Gyrus BA22 |
|  |  | 52 | -20 | 0 | 2E-02 | 7E-05 | 3.8 | R Superior Temporal Gyrus BA22 |
| 3 | 920 | -58 | -46 | 16 | 2E-02 | 2E-06 | 4.6 | L Superior Temporal Gyrus BA22 |
|  |  | -56 | -36 | 22 | 2E-02 | 9E-05 | 3.8 | L Insula BA13 |
| NM>M: 144 foci, 12 experiments, 286 subjects |  |  |  |  |  |  |  |  |
| 4 | 912 | -50 | -36 | 56 | 1E-02 | 2E-05 | 4.1 | L Inferior Parietal Lobule BA40 |
|  |  | -46 | -40 | 52 | 1E-02 | 4E-05 | 4.0 | L Inferior Parietal Lobule BA40 |
| 5 | 736 | -48 | 8 | 36 | 2E-02 | 3E-06 | 4.5 | L Precentral Gyrus BA6 |

ALE, anatomic likelihood estimation; M, musicians; NM, non-musicians; GM, grey matter; WM, white matter; BA, Brodmann area; P, p-value; Z, peak z-value; R, right; L, left; NA, not enough available observations.

#### 2.3.2. Functional studies

The functional ALE meta-analysis included 46 experiments and 1265 participants. The contrast M > NM resulted in large and significant peak clusters of the bilateral superior temporal gyrus (the bilateral primary auditory cortices) extending to the left insula, the left inferior frontal gyrus, and the left precentral gyrus (primary motor cortex, M1). The comparison NM > M resulted in smaller peak clusters of the left inferior parietal lobule and the left precentral gyrus (**Figure 1**, **Table 2**).

### 2.4. Meta-analytic connectivity modelling (MACM)

MACM was performed to functionally segregate the behavioural contribution and the patterns of coactivation of each music-related region-of-interest (ROI) resulted from the structural (n=5) and functional (n=5) ALE meta-analyses. Five-millimetre ROIs were created using Mango^36^ and imported into the BrainMap^37^ database separately using Sleuth^38^. Foci from each identified study were extracted and a secondary GingerALE meta-analysis was performed aiming to identify the functional network of each ROI, namely its functional connectivity (FC)^39^ (**Supplementary Table 4**). Finally, the functional characterization of each ROI was described using Sleuth and focused on behavioural domains (e.g., action, perception, emotion, cognition and interoception) and paradigm classes (e.g., pitch discrimination, finger tapping, music comprehension, go/no-go) (**Supplementary Table 5**).

#### 2.4.1. Structural ROIs

The right superior temporal gyrus ROI (**Figure 2a**) showed co-activation with left superior temporal gyrus, right precentral gyrus, left medial frontal gyrus, left cerebellum, and left thalamus. Relevant behavioural domains within its boundaries include execution, speech, memory, music, emotion, reward, and auditory perception; and experimental paradigms including emotion induction, finger tapping, music comprehension and production, passive listening, reasoning/problem solving, and phonological, pitch, semantic, syntactic, and tone discrimination.

**Figure 2.**
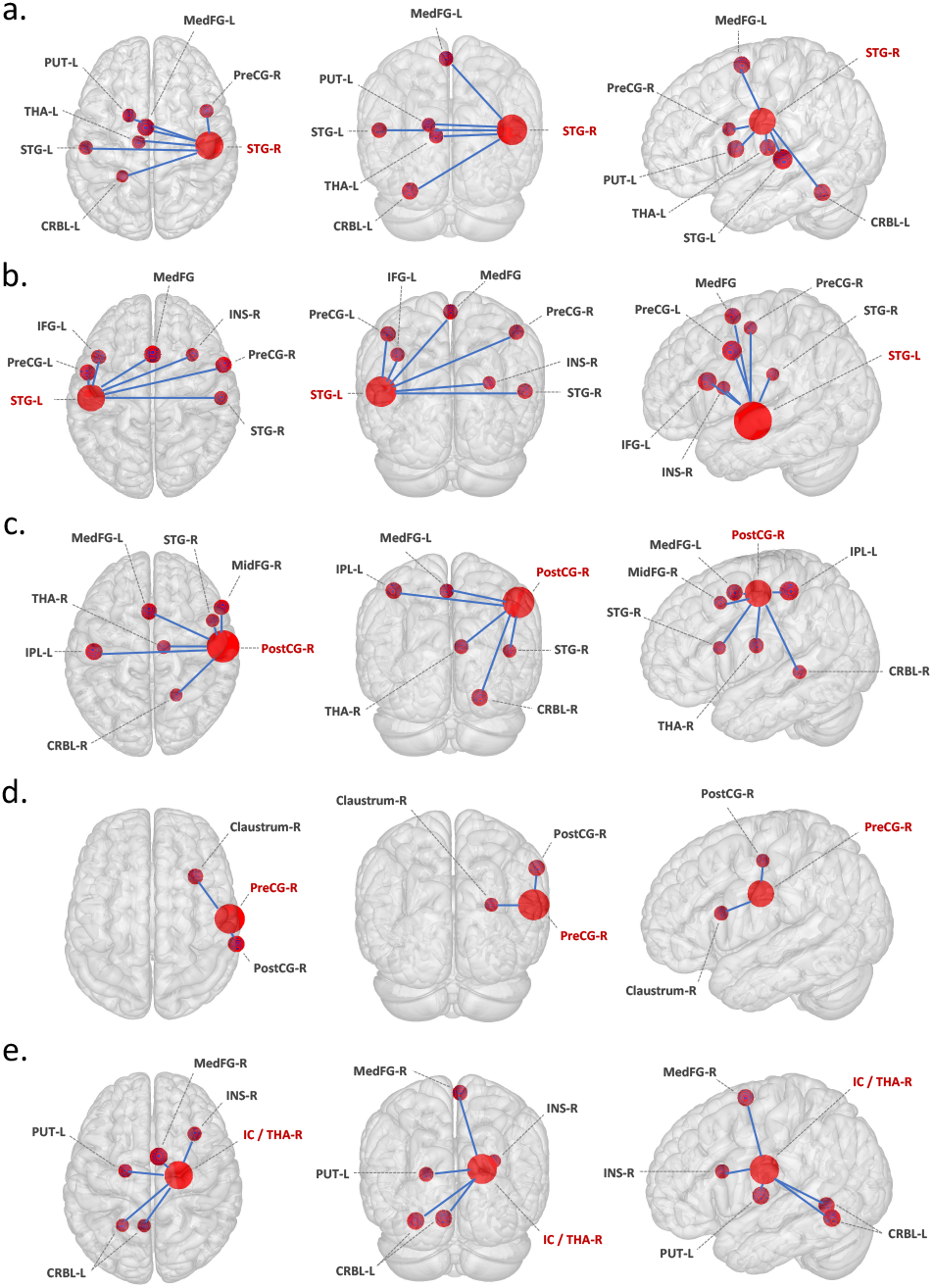
Meta-analytic connectivity modelling of regions-of-interest that resulted from the structural ALE meta-analysis, at cluster level inference p < 0.05 (FWE). ROIs, music-related regions-of-interest; P, p-value; Z, peak z-value; R, right; L, left. **ROIs**: a. STG-R, right superior temporal gyrus; b. STG-L, left superior temporal gyrus; c. PostCG-R, right postcentral gyrus; d. PreCG-R, right precentral gyrus; e. IC/THA-R, internal capsule (including right thalamus). **Co-activated areas**: claustrum; CRBL, cerebellum; IFG, inferior frontal gyrus; IPL, inferior parietal lobule; INS, insula; MedFG, medial frontal gyrus (pre-motor); MidFG, middle frontal gyrus (pre-frontal); PostCG, postcentral gyrus (primary somatosensory cortex or S1); PreCG, precentral gyrus (primary motor cortex or M1); PUT, putamen; STG, superior temporal gyrus (primary auditory cortex), THA, thalamus. To conduct MACM, music-related ROIs were created in Mango (http://rii.uthscsa.edu/mango//userguide.html) with a 5mm-radius sphere. For visualization purposes, the music-related ROI radius was increased to 10mm, while co-activated areas were created with a 5mm-radius sphere. Last search in Sleuth, 10.10.2021 (http://www.brainmap.org/sleuth/).

The left superior temporal gyrus ROI (**Figure 2b**) showed co-activation with right superior temporal gyrus, right insula, right inferior frontal gyrus, bilateral precentral gyrus, and left medial frontal gyrus. Relevant behavioural domains within its boundaries include execution, speech, motor learning, attention, language, speech, memory, music, emotion, and auditory perception; and experimental paradigms including emotion induction, finger tapping, music comprehension and production, passive listening, reasoning/problem solving, visuospatial attention, and oddball, orthographic, phonological, pitch, semantic, and tone discrimination.

The right postcentral gyrus ROI (**Figure 2c**) showed co-activation with left medial frontal gyrus, left parietal lobule, right middle frontal gyrus, right thalamus, right superior temporal gyrus, and right cerebellum. Relevant behavioural domains within its boundaries include execution, motor learning, attention, respiration regulation, and auditory perception; and experimental paradigms including finger tapping, passive listening, visuospatial attention, and oddball, tactile, and tone discrimination.

The right precentral gyrus ROI (**Figure 2d**) showed co-activation with claustrum and insula. Relevant behavioural domains within its boundaries include execution, attention, speech, temporal processing, and emotional processing; and experimental paradigms including finger tapping and visuospatial attention.

The right internal capsule ROI (**Figure 2e**), including the right thalamus as the nearest grey matter, showed co-activation with left thalamus, right medial frontal gyrus, right insula, and left cerebellum. Relevant behavioural domains within its boundaries include execution, speech, attention, memory, reasoning, emotion, reward, and auditory perception; and experimental paradigms including emotion induction, finger tapping, passive listening, reward, and tone discrimination.

#### 2.4.2. Functional ROIs

The left inferior frontal gyrus ROI (**Figure 3a**) showed co-activation with right inferior frontal gyrus, left inferior and superior parietal lobules, left medial frontal gyrus, left fusiform gyrus, and right caudate. Relevant behavioural domains within its boundaries include execution, speech, attention, language, memory, music, reasoning, social cognition, emotion, and auditory perception; and experimental paradigms including encoding, finger tapping, music comprehension and production, passive listening, reasoning/problem solving, reward, and phonological, semantic, tactile, and tone discrimination.

**Figure 3.**
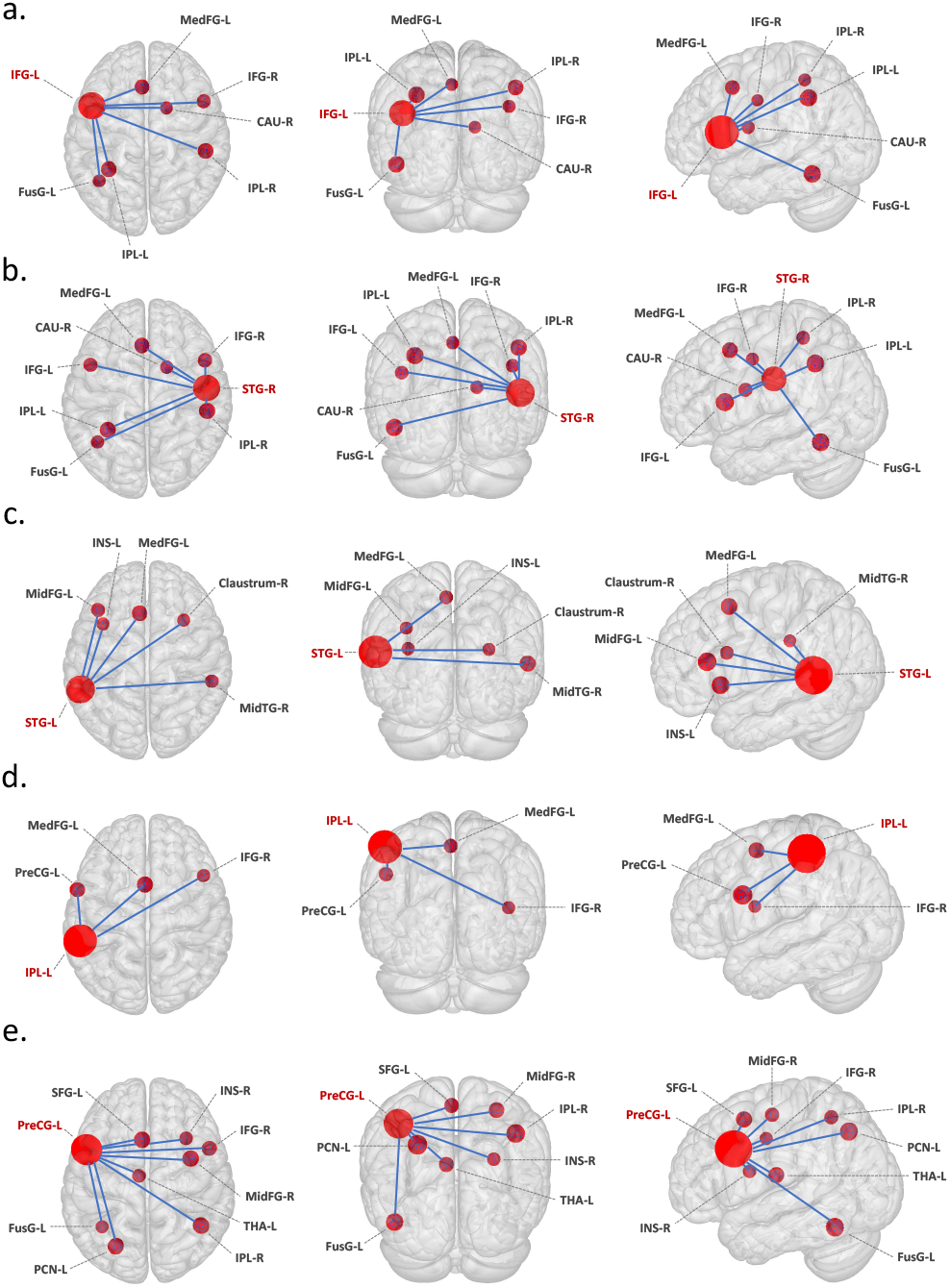
Meta-analytic connectivity modelling of regions-of-interest that resulted from the functional ALE meta-analysis, at cluster level inference p < 0.05 (FWE). ROIs, music-related regions-of-interest; P, p-value; Z, peak z-value; R, right; L, left. **ROIs**: a. IFG-L, left inferior frontal gyrus; b. STG-R, right superior temporal gyrus; c. STG-L, left superior temporal gyrus; d. IPL-L, left inferior parietal lobule; e. PreCG-R, right precentral gyrus. **Co-activated areas**: CAU, caudate; claustrum; CRBL, cerebellum; FusG, fusiform gyrus; IFG, inferior frontal gyrus; IPL, inferior parietal lobule; INS, insula; MedFG, medial frontal gyrus (pre-motor); MidFG, middle frontal gyrus (pre-frontal); PostCG, postcentral gyrus (primary somatosensory cortex or S1); PreCG, precentral gyrus (primary motor cortex or M1); PCN, precuneus; PUT, putamen; SFG, superior frontal gyrus; STG, superior temporal gyrus (primary auditory cortex), THA, thalamus. To conduct MACM, music-related ROIs were created in Mango (http://rii.uthscsa.edu/mango//userguide.html) with a 5mm-radius sphere. For visualization purposes, the music-related ROI radius was increased to 10mm, while co-activated areas were created with a 5mm-radius sphere. Last search in Sleuth, 10.10.2021 (http://www.brainmap.org/sleuth/).

The right superior temporal gyrus ROI (**Figure 3b**) showed co-activation with bilateral inferior frontal gyrus, bilateral inferior parietal lobule, left medial frontal gyrus, left fusiform gyrus, and caudate. Relevant behavioural domains within its boundaries include execution, speech, attention, language, memory, music, reasoning, social cognition, emotions, and auditory perception; and experimental paradigms including finger tapping, music comprehension and production, passive listening, reasoning/problem solving, reward, theory of mind, and phonological, semantic, tactile, and tone discrimination.

The left superior temporal gyrus ROI (**Figure 3c**) showed co-activation with right superior temporal gyrus, right middle temporal gyrus, right claustrum, left insula, and left medial frontal gyrus. Relevant behavioural domains within its boundaries include execution, speech, attention, language, memory, reasoning, social cognition, emotion, and auditory perception; and experimental paradigms including divided auditory attention, emotion induction, emotional body language perception, encoding, finger tapping, music comprehension and production, reasoning/problem solving, theory of mind, visuospatial attention, and oddball, phonological, pitch, semantic, and tone discrimination.

The left inferior parietal lobule ROI (**Figure 3d**) showed co-activation with left medial frontal gyrus, right inferior frontal gyrus, and left precentral gyrus. Relevant behavioural domains within its boundaries include execution, speech, motor learning, attention, memory, music, reasoning, social cognition, emotion, and auditory perception; and experimental paradigms including emotion induction, finger tapping, motor learning, reasoning/problem solving, reward, visuospatial attention, and phonological, semantic, tactile and tone discrimination.

The left precentral gyrus ROI (**Figure 3e**) showed co-activation with left precuneus, left superior frontal gyrus, right inferior frontal gyrus, right inferior parietal lobule, right claustrum, left fusiform gyrus, left thalamus, and right middle frontal gyrus. Relevant behavioural domains within its boundaries include execution, speech, motor learning, attention, language, memory, music, reasoning, social cognition, temporal processing, emotion, sleep, and auditory perception; and experimental paradigms including divided auditory attention, emotion induction, encoding, finger tapping, music comprehension, reasoning/problem solving, reward, theory of mind, visuospatial attention, and oddball, phonological, pitch, semantic, tactile, and tone discrimination.

## 3. Discussion

The link between musical expertise and humans’ cognitive functions has been explored with great interest since the times of Pythagoras. Recent years reveal a renewed and more than vivid attention to the topic, as reflected in the rising number of empirical research in the past half-century^40,41^. Decades of investigations in psychology, cognitive and translational neuroscience have attempted to foster our understanding of the neurocognitive processes underlying musical expertise. Thus, longterm musical training has been associated with neuro-anatomical and -functional specializations in brain regions engaged in multimodal (audio-visual) sensory and sensorimotor perception, integration and predictions as well as fine movement control^15,16^. Furthermore, the duration and intensity of training has been associated with improvements in general cognition, ranging from working memory, intelligence, executive functions and inhibitory control^5,8–10^. However, as mentioned before, this rapidly growing field of research is also characterized by some methodological inconsistencies (e.g., sample differences and neglected background variables), and sometimes shows discrepant results and controversial interpretations of the findings. Such limitations, alongside with the absence of a metaanalysis, has prevented the plethora of variegated findings to ultimately converge into a unified picture of the neuroanatomy of musical expertise.

To address this lack in the literature, we performed a comprehensive and quantitative meta-analysis of neuro-anatomical and -functional studies investigating brain changes associated with long-term musical training. Our coordinate-based anatomic/activation likelihood estimation (ALE) meta-analysis effectively summarizes decades of research in the field and finally provides a consistent and controversies-free picture of the core brain regions engaged in and influenced by long-term music processing and production. To better characterize the emergent neural network of musical expertise, we performed meta-analytic connectivity modelling analyses (MACM) and functionally linked each node of the music network to specific cognitive functions. By discussing the main results of the metaanalysis alongside with the observations derived from MACM, we ultimately provide a comprehensive view of the anatomical, functional, and cognitive substrates of musical expertise. This discussion is organized in three main paragraphs: ‘the ear’, ‘the body’ and ‘the heart’, elaborating on the emergent fronto-temporal, sensorimotor and interoceptive networks respectively. Notably, MACM further allows us to strengthen the notion that musical training represents a stimulating multisensory experience engaging not only sensory and motor functions strictly related to acoustic and motor processes, but a wide variety of high-order cognitive functions from working memory, attention, executive functions and emotional regulation^1,10,15,16,42^. To conclude, we argue that the observed music-related neuroanatomical and -functional changes represent an interface between *nature* and *nurture* effects. Namely, gene-environment interactions and other background variables likely interacted with brain maturation processes ultimately influencing the neuroplasticity mechanisms responsible for the observed training-specific neuroanatomical and -functional changes.

### 3.1. Characteristics of the included studies

The publications included in this systematic review and meta-analysis reported a clear research question, inclusion and exclusion criteria for participants, description of methods and explicit results. Most of the studies used state-of-the-art techniques and computational MRI-tools, important for the support of standardization and reproducibility of neuroimaging studies. However, some of the studies lacked important demographic data such as the years of education, age of musical training onset, and current time of musical practice, which may influence behavioural tasks and neuroimaging data. Thus, our research encourages to adopt in future studies standardized tools specifically designed and validated for assessing musical expertise^43^.

### 3.2. Structural and functional neuroplasticity in musical expertise

Our results highlight that expert musicians exhibited higher GM volume in the bilateral superior temporal gyri and right postcentral gyrus and greater WM volume in the right internal capsule bundle and corticospinal tract, as compared to non-musicians. Additionally, musicians exhibited higher activity of the bilateral superior temporal gyri, left inferior frontal gyrus, left precentral gyrus, and left insula. On the other hand, musicians had lower GM volume in areas of the sensorimotor cortex and no WM structure was found to have larger volume in non-musicians as compared to musicians. Finally, musicians exhibited lower neurofunctional activation of the inferior parietal lobule and motor cortex during a variety of cognitive tasks.

#### The ear: Enhanced frontotemporal auditory network in musicians

One of our main findings shows enlargement of GM volume in musicians located in medial and posterior superior temporal regions, with clusters extending into primary and secondary auditory cortices. These regions include neuronal assemblies dedicated to encoding of spectro-temporal features of sounds relevant to music^44^, such as the discrete pitches forming the Western chromatic scale and fine changes in pitch intervals^45^. More specifically, it seems that the posterior supratemporal regions are more involved in encoding the height of pitch, whereas the anterior regions are representing the chroma, that is the pitch category irrespectively of the octave^46^. Moreover, these areas participate in auditory imagery of melodies^47^ and in the processing of the contour and Gestalt patterns of melodies, allowing for recognition and discrimination of mistakes^48^.

Beyond music-related functions, functional characterization analyses of our ROIs (**Supplementary Table 5**) show that superior temporal regions are usually recruited for phonological processing and multimodal integration of sensory information. Accumulating evidence has shown that the superior temporal sulcus and posterior superior temporal gyrus, together with early auditory regions (HG), are involved for the processing of speech sounds, abstract representation of speech sounds, as well as more general language, phonology and sematic processing and audio-visual integration. Therefore, temporal regions seem to represent fundamental structures for both language and music processing^49^. MACM further revealed that auditory cortices tend to co-activate with insula, (pre)motor regions, inferior and medial frontal gyri, thalamus and cerebellum, confirming the relevance of extended cortico-subcortical audio-motor coupling for rhythm processing in language and music ^50–53^. Further supporting this view, MACM showed that the inferior frontal gyrus co-activates with motor areas in the cortex and the basal ganglia (see next paragraph, ‘The body’), and with parietal areas related to the dorsal auditory pathway.

The inferior frontal gyrus has been described as an important hub of both the dorsal and ventral auditory streams. The dorsal auditory stream connects the auditory cortex with the parietal lobe, which projects in turn to the inferior frontal gyrus pars opercularis (Brodmann area 44). The inferior frontal gyrus has been related to the articulatory network, dedicated to specific functions of speech comprehension and production, and highly connected to premotor and insular cortices^54^. The ventral auditory stream connects the auditory cortex with the middle temporal gyrus and temporal pole, which in turn connects to the inferior frontal gyrus pars triangularis (Brodmann area 45). This area has been associated with semantic processing^55^. These two regions within the inferior frontal gyrus constitute Broca’s area. The supramarginal gyrus is also a relay of the dorsal auditory stream involved in processing of complex sounds, including language and music^56^. As such, it is considered an integration hub of somatosensory input^57^.

The parietal lobe has been also described as an integration area of sensory inputs. The superior parietal lobule includes Brodmann areas 5 and 7, which are involved in somatosensory processing and visuomotor coordination, respectively. The inferior parietal lobule includes Brodmann areas 39 and 40, the angular gyrus and supramarginal gyrus, respectively. The angular gyrus has been related to projection of visual information to Wernicke’s area, memory retrieval and theory of mind^58^. MACM revealed that the parietal lobe co-activates with sensorimotor cortices and the inferior frontal gyrus.

#### The body: Enhanced sensorimotor functions in musicians

The precentral and postcentral gyri represent the primary motor and somatosensory cortex, respectively. These two areas are divided by the central sulcus, whose extension represent the sensation and motion of segregated body parts. Our findings show both convergent and divergent effect of musical training in these areas, suggesting a more complex picture than previously thought. For example, neuroadaptations in the sensorimotor system may vary depending on the musical instrument of use^59^. MACM revealed that the primary motor cortex co-activates with an extensive network that includes the frontal pole, limbic areas such as the anterior cingulate cortex and insula, and parietal areas such as the precuneus. It also revealed that the primary somatosensory cortex coactivates with motor and pre-motor areas, basal ganglia, thalamus, and the cerebellum. A dedicated temporal processing network has been described by Kotz and Schwartze^51^ including such areas, which are important for implementing sequential actions, as well as to form predictions about the timing of external events. Healthy motor performance relies on a functional loop established by the basal ganglia and supplementary motor area that maintains adequate preparation for sequential movements. The supplementary motor area prepares for predictable forthcoming movements, keeping the system “ready”. Once the movement starts, the supplementary motor area’s readiness activity stops. This cycle engages with BG discharges after each sub-movement within an automatized sequence^60^. The loop requires an internal cue to coordinate the cycle.

The basal ganglia are nuclei of neurons important for the initiation and suppression of movements. In the motor loop of the basal ganglia (BG), inputs from motor cortices project to the dorsal striatum, composed by the putamen and caudate. In the presence of adequate dopaminergic signalling, the ‘direct pathway’ (cortex – striatum – internal pallidum – thalamus – cortex) works to facilitate movement, while the ‘indirect pathway’ suppresses it (cortex – striatum – external pallidum – subthalamic nucleus – thalamus – cortex). Zooming into disinhibition processes, the striatum transiently inhibits the pallidum, and in turn, the motor area of the thalamus is disinhibited and is free to project back to the motor cortex, initiating a motor program that flows down the corticospinal tract. Similarly, the subthalamic nucleus in the indirect pathway is transiently inhibited when suppressing movement, increasing the inhibition of the pallidum over the thalamus, therefore blocking the motor cortex activity^61^. Our findings show neuroadaptive processes in the putamen and caudate of musicians (striatum), presumably reflecting effective disinhibition mechanisms as seen by fine movement control.

The cerebellum has been shown to play a crucial role in multiple cognitive processes such as sensory discrimination, rhythmic perception and production, working memory, language, and cognition^62^. Previous fMRI studies in humans suggest that the cerebellum shows segregated activations for motor and cognitive tasks. Motor tasks seem to activate lobules IV-VI in the superior parts of the anterior cerebellum. In contrast, attentional or working memory tasks activate posterior cerebellar hemispheres, namely lobule VIIA, which is divided to crus I and crus II, as well as lobule VIIB^63^. Musicians and non-musicians show GM volume differences in the cerebellum, specifically in area Crus I. In our study, this area did not survive correction for multiple comparisons, however MACM revealed that the cerebellum is functionally connected to auditory cortices, somatosensory cortices, and the thalamus. It has been demonstrated that the activity in crus I/II has a specific relationship with cognitive performance and is linked with lateral prefrontal areas activated by cognitive load increase^64^. In other words, the crus I/II seems to optimize the response time when the cognitive load increases. Additionally, it has been suggested that crus I/II is associated with beat discrimination thresholds. Thus, there is a positive correlation between GM volume in crus I and beat discrimination performance, evidenced by enhanced ability in musicians^65^.

#### The heart: Enhanced interoceptive areas in musicians

Among the other results, our meta-analysis reported higher functional activation of left insula in musicians as compared to non-musicians. MACM analyses reported the left insula in a functional network that connects inferior frontal gyrus with precentral gyrus, middle frontal gyrus and parietal lobule bilaterally (**Supplementary Table 4**).

It has been proposed that the insula and the anterior cingulate cortex (ACC) are part of the salience network, and coordinate interactions between the default-mode network and the central executive network^66^. The ACC has been related to cognitive and emotional processing. The cognitive component projects to prefrontal, motor, and parietal areas to process top-down and bottom-up stimuli. The emotional component features connections from the insula to amygdala, nucleus accumbens, hypothalamus and hippocampus, with the scope to assess the salience of emotional and motivational information^67^. Moreover, the insula integrates information from the internal physiological state, and projects to the ACC, ventral striatum and prefrontal cortex to initiate adaptive responses^68^. Thus, enhanced function of these areas after musical training may be associated with a more efficient coordination between interoceptive, emotional, salience and central executive networks.

#### White matter

M exhibited larger clusters of WM as compared to NM in the internal capsule and cortico-spinal tract. While previously thought to be rather passive tissues, WM tracts are now consistently associated with an active modulatory role in information flow between brain regions^61^. Indeed, myelin regulates the speed of action potential transfer within and between GM structures and further provide metabolic support to local neural cells. WM changes are commonly observed during learning and associated with fast, accurate and coordinated motor sequences^24^.

The internal capsule is a WM structure which connects basal ganglia regions and carries information from and to surrounding cerebral cortex. Connecting fibres in basal ganglia might be thickened by musical expertise because of their involvement in motor control, rhythmic processing, sequence learning, reinforcement learning and memory processes^69^. In general, basal ganglia structures are recruited during working memory processing for musical motifs^70^ and the most ventral regions are a core structure of the reward circuit. Interestingly, they are found to be more active in musicians as compared to non-musicians while listening to expressive music^71^.

The corticospinal tract allows the motor plans originated in the cortex to be transferred to motor nuclei in the spinal cord and to finally regulate the activity of muscle effectors. The myelination and integrity of the corticospinal tract has been observed to be increased in expert musicians ^24,25^, and is further influenced by the time of onset of musical practice ^28^ with early onset musician showing the greatest diffusivity.

### 3.3. (Dis)similarities between anatomical and functional studies

The meta-analyses on neuroanatomical and -functional changes coherently show greater GM volumes and increased functional engagement of superior temporal gyrus bilaterally, together with pre- and post-central gyri in expert musicians as compared to laypersons. Functional studies further agree on the pivotal involvement of left inferior frontal gyrus (BA9, BA44) next to superior temporal gyrus bilaterally in musicians. However, dissimilarities emerge when looking at pre- and post-central regions: right precentral gyrus (right primary motor cortex (M1)) is reduced in musicians, while the right postcentral gyrus (right primary somatosensory cortex (S1)) is increased. Functional studies show, instead, that there is increased activity in left precentral gyrus in the inferior frontal gyrus - superior temporal gyrus - insula network, and reduced activity of the left precentral gyrus in a cluster which extends into the left parietal lobule.

While results pertaining to the frontotemporal auditory network and the sensorimotor network have been discussed in ‘*The ear*’ and ‘The body’ paragraphs above, we here speculate that the enlargement of S1 in musicians is associated with a more sophisticated representation of the sensorimotor periphery^17^ and that the increased left inferior frontal gyrus - precentral gyrus - superior temporal gyrus -insula activation at the expense of the M1-parietal lobule network may be related to the acquisition of accurate and automatized motor programs in musicians^3^. In agreement with early studies, we lastly argue that the hemispheric asymmetry may be related to the music instrument played and the dominant hand of the musicians^72^, but interhemispheric transfer effects are possible with motor sequence learning ^73^. However, longitudinal studies should further elucidate on the heterogeneity of structural and functional adaptations associated with intensive and long-lasting motor training.

### 3.4. Limitations and future perspectives

This comprehensive review and meta-analysis had the scope to summarize decades of research investigating neuro-anatomical and -functional changes associated with musical expertise. Our qualitative review highlights that previous studies in this field are characterized by heterogeneity of methods, paradigms, and sample backgrounds, as well as relevant missing information. While arguing that the field will benefit from more clarity (e.g., thorough description of methods) and consistency, we also delineate limitations for our meta-analysis. For example, we set a contrast based on the comparison M vs NM with the aim to narrow down the heterogeneity of the sample and methods in use. However, by doing so we relied on two assumptions: (1) the data we pool is based on best research practices; (2) the validity of the GingerALE method. Indeed, to conduct the ALE meta-analysis, we pooled peak coordinates derived from the included studies, rather than using original raw structural MRI images. Thus, the accuracy of our findings relies on the result of a statistical estimation of coordinate-based anatomic foci (input), treated as spatial probability distributions centred at the given coordinates. The heterogeneity of the methods in use in previous studies (ranging from preprocessing software, smoothing, statistical thresholds and participants’ characteristics) are not under our control and represent potential confounders for the results. Perhaps a regression-based assessment of the influence of those heterogenous factors on the findings would sharpen the results. However, meta-regression analysis is not compatible with GingerALE. When assessing publication bias using the Fail Safe-N analysis, we found adequate robustness of our results, with only 2 ROIs showing an FSN below of the minimum imposed in each of the ALE within contrasts (BA2, BA4 in the structural ALE and BA22, BA6 in the functional ALE), thus, indicating an overall robust convergence of foci our study (further information is reported in **Supplementary Table 6**).

Lastly, on a more theoretical perspective, our results contribute but do not solve the long-standing “nature vs nurture” debate. Indeed, based on evidence that musical training stimulates higher-cognitive functions, auditory-motor integration, attention, memory and engages reward networks, some have suggested that it may be particularly effective in driving neuroplastic mechanisms^74^. However, we are indeed blind to whether the highlighted differences emerging when comparing M vs NM are training-dependent or due to innate predispositions. Altogether, the most reasonable conclusion is that the observed neuro-anatomical and -functional changes may be attributed to the interaction between brain maturation processes and gene-environment interactions ^13,77,50^. Notably, multiple studies demonstrated a strong correlational link between the length of musical training and neuroanatomical and -functional changes. For instance, the study conducted by Gaser & Schlaug^77^ reported that amateur musicians showed an intermediate increase in gray matter volume when compared to NM and M, supporting the idea of use-dependent structural changes. The same pattern was found when comparing cognitive abilities, with amateurs showing higher cognitive abilities than NM, but lower than M^78^. To be noted, however, this research field suffers of the paucity of longitudinal (f)MRI studies conducted with children, which thus far amount only to seven^3–7,79,80^, next to one 15-week long study in adults ^81^. Longitudinal studies are the only ones promising to better elucidate on the causal link between musical training and neural adaptations. Our work, on the other hand, pools a large quantity of anatomical and functional MRI studies conducted over >20 years of world-wide research. By doing so, it bears the potential to achieve an unprecedented signal-to-noise ratio, so to filter out the mediating influence of background, psychological and other environmental factors, and to effectively isolate music-related neuroplastic changes. Thus, we here provide, within the delineated limits, a consistent view of the neuroanatomy of neural expertise. We hope our work would better inform future basic and comparative research in the field of auditory and cognitive neuroscience and that we encouraged translational approaches bridging to the clinical field ^82,83^.

## 4. Conclusions

The neuroanatomical and functional changes in observed in the musician’s brain have been repeatedly regarded as the ideal scenario to investigate neuroplastic mechanisms. Yet, decades of research in cognitive neuroscience have provided a scattered and partially controversial series of findings. The present coordinate-based meta-analysis represents a comprehensive and quantitative attempt to summarize existing literature and provide a unified picture of the neuroanatomy of musical expertise. We show that music experts exhibit bilateral cortico-subcortical neuroanatomical and -functional differences as compared to laypersons. This comprehensive work strengthens the view that musical training represents a beneficial and stimulating multisensory experience which engages a wide variety of neurocognitive functions.

## 5. Methods

### 5.1. Literature search, screening, and extraction

This systematic review and meta-analysis followed procedures from the Cochrane Handbook for Systematic Reviews^84^ and from the Centre for Reviews and Dissemination (Centre for Reviews and Dissemination, 2014). The review protocol was registered with PROSPERO No. [<u>CRD42017060365</u>]. This review was carried in accordance with the PRISMA statement^85^.

Systematic search was performed using PubMed, PsycInfo and Scopus, of publications that reported brain structural or functional differences between M and NM. The search (March 2021) included MeSH terms (“music”, “education”, “brain”, “motor skills”, “magnetic resonance imaging”) and key words (“musical training”, “musician”). No years or places of publication were imposed.

For qualitative synthesis, studies were included if they met the following criteria: (1) studies comparing brain structure and function between musicians and non-musicians, (2) in adult population, (3) by means of magnetic resonance imaging, in either structural modality (e.g., voxel-based morphometry [VBM]) or functional modality (e.g., functional magnetic resonance imaging[fMRI]). For the final quantitative synthesis (meta-analysis), studies were included only if the results were reported in stereotactic coordinates either Talairach or Montreal Neurological Institute (MNI) threedimensional-coordinate system.

Studies were excluded using the following criteria: (1) review articles with no original experimental data, (2) neuroimaging data from non-MRI studies (e.g., PET), (3) pathological population, (4) longitudinal designs, (5) functional connectivity analyses, and (6) analyses based on region-of-interest (ROI) rather than whole-brain (only quantitative synthesis).

Two reviewers (AC and VP) independently screened by title and abstract and selected articles for fulltext review and performed full-text reviews. Screening and data extraction were performed using the Covidence tool^86^. Any disagreements that arose between the reviewers were resolved through discussion or by a third and/or fourth reviewer (LB, EB).

From each study, the following variables were extracted: first author, year of publication, population of interest, number of participants, age, sex, absolute pitch, musical feature, years of education, years of musical training, age of musical training onset, weekly training, musical instrument, MRI-system, MRI-model, head-coil, image acquisition parameters of T1, T2* and DWI sequences, repetition time (TR), echo time (TE), voxel size, analysis method and software. The main outcome to extract was any difference in structure or function, in stereotactic coordinates, comparing a musician group and a nonmusician group. If any of these points were not reported in the original article, authors were contacted to retrieve this information. Six authors were contacted, with 2 positive answers.

### 5.2. Quality assessment of MRI studies

Criteria for MRI quality reporting was selected from a set of guidelines for the standardized reporting of MRI studies^33,34^. Such guidelines dictate a more consistent and coherent policy for the reporting of MRI methods to ensure that methods can be understood and replicated.

### 5.3. Activation likelihood estimation (ALE)

To test the convergence of findings from the neuroimaging studies, we used the anatomic/activation likelihood estimation (ALE) method implemented in the GingerALE software v3.0.2^32^, a widely used technique for coordinate-based meta-analysis of neuroimaging data. Statistically significant foci from between-group contrasts were extracted and recorded for each study. If necessary, coordinates were converted from Talairach coordinates to MNI space using the Lancaster transform (icbm2tal) incorporated in GingerALE^35,87^. The ALE method uses activation foci (input) not as single points, but as spatial probability distributions centred at the given coordinates. Therefore, the algorithm tests to what extent the spatial locations of the foci correlate across independently conducted MRI studies investigating the same construct and assesses them against a null distribution of random spatial association between experiments^46^. Statistical significance of the ALE scores was determined by a permutation test using cluster-level inference at p < 0.05 (FWE), with a cluster-forming threshold set at p < 0.001. First, we used the ALE meta-analytic technique to identify brain differences measured by MRI between musicians (M) and non-musicians (NM) with the aim of comprehensively examine the neural signatures of musical expertise. Two independent ALE meta-analyses were conducted for structural studies and functional studies. To test the directionality of the M vs NM contrast, foci were pooled reporting higher volume/activity in musicians (M > NM) and lower volume/activity in musicians (NM > M) for both structural and functional studies.

### 5.4. Meta-analytic connectivity modelling (MACM)

Meta-analytic connectivity modelling (MACM) was performed to analyse co-activation patterns of music-related regions-of-interest (ROI) resulted from the structural (n=5) and functional (n=5) ALE meta-analyses, independently, and to functionally segregate each region’s putative contribution to behavioural domains and paradigm classes according to the BrainMap platform^36,37,88^.

Large-scale databases such as BrainMap store results obtained from published brain activation (functional) and brain structure (voxel-based morphometry) studies^89,90^. Such databases can be taken into advantage with a meta-analytic approach focusing on the co-activation of brain regions with a specific ROI across all kinds of different mental processes, rather than to a specific mental process. Thus, MACM identifies the functional network of the ROI, namely, its functional connectivity (FC). Traditionally, in fMRI studies, two brain regions are functionally connected when there is a statistical relationship between the measures of neuronal activity, by means of the blood-oxygen-level-dependent signal (BOLD), both during resting-state (task-free FC) or performing a specific task (task-dependent FC). In contrast, MACM relies on patterns of co-activation across many different tasks and allows to examine task-based FC in a general manner^88,91,92^. Thus, MACM provides a data-driven and unbiased approach to determine the connectivity “signature” of a given ROI.

Co-activation analyses were performed using Sleuth^37^ and GingerALE^32^ from the BrainMap platform. To identify regions of significant convergence, an ALE meta-analysis was performed over all foci retrieved after searching Sleuth by each music-related ROI independently and included the experiment level search criteria of “context: normal mapping” and “activations: activation only”. Music-related ROIs were created in Mango^93^ with a 5mm-radius sphere. The results of each ROI search were exported to GingerALE, and a permutation test was conducted using cluster-level inference at p < 0.05 (FWE), with a cluster-forming threshold set at p < 0.001.

Finally, MACM allows to conduct functional profiling of ROIs to study brain-behaviour relationships at a meta-analytic level. In other words, through the BrainMap platform, it is possible to objectively characterize a given ROI in terms of its cognitive/behavioural function which are based on the meta-data that is stored in the database^39^. Thus, tasks in the database are coded in a way that is possible to conduct a behavioural profile of ROIs that resulted from an ALE meta-analysis. The tasks are coded in two dimensions: behavioural domains (BD) and paradigm classes (PC). As the present study has two independent meta-analyses, one for structural studies and one for functional studies, MACM was divided into ROIs that resulted from the structural ALE meta-analysis and ROIs that resulted from the functional ALE meta-analysis. The functional characterization of music-related ROIs was based on the BD meta-data categories available for each neuroimaging study in the database which include action, perception, emotion, cognition and interoception. PC refer to paradigms that have been used repeatedly by different researchers with only minor changes. Such paradigms have become widely known and accepted by the neuroimaging field (e.g., pitch discrimination, finger tapping, music comprehension, go/no-go). A BD refers to the categories and sub-categories of mental operations likely to be isolated by the experimental contrast; a PC is the experimental task isolated by the contrast of interest. Notably, multiple BDs and PCs may apply for a given experiment^94^.

All meta-analytic results (ALE maps) were visualized using Mango^36^ on the MNI152 1mm standard brain, and resulting coordinates were cross-referenced to the Harvard-Oxford Cortical and Subcortical Atlas and the Juelich Histological Atlas via NeuroVault^95^ and FSLeyes^96^, respectively.

### 5.5 Fail-Safe N analysis (FSN)

As all meta-analyses, coordinate-based meta-analyses such as ALE can be subject to different forms of publication bias which may impact results and invalidate findings (e.g., the “file drawer problem”). Thus, the Fail-Safe N analysis (FSN)^97^ was performed as a measure of robustness against potential publication bias. It refers to the amount of contra-evidence that can be added to a meta-analysis before the results change and can be obtained for each cluster that survives thresholding in an ALE meta-analysis. For normal human brain mapping, it is estimated that a 95% confidence interval for the number of studies that report no local maxima varies from 5 to 30 per 100 published studies. Therefore, the minimum FSN was defined as 30% of total studies for each CBMA. A higher FSN indicates more stable results and hence a higher robustness.

## Acknowledgments

The Center for Music in the Brain (MIB) is supported by the Danish National Research Foundation (grant number DNRF 117). The authors wish to thank Hella Kastbjerg for assistance with language check and proof reading. Leonardo Bonetti is supported by the Carlsberg Foundation (CF20-0239) and Linacre College, University of Oxford.

## Authors’ contributions

AC, VPN and EB designed the meta-analysis. VPN guided AC in the initial steps of the meta-analysis after which AC and VPN conducted the screening of the studies, and EB and LB controlled the final selection. AC performed the initial analyses, wrote the first draft of the manuscript, and prepared the initial versions of the tables and figures. VPN performed analyses, prepared the figures and tables and edited paragraphs in the Methods, Results and Discussion sections. All authors contributed to the final version of the manuscript. PV contributed to financially support the study.

## Data availability

The data supporting the findings of this study is freely available at the Open Science Framework (OSF) website https://osf.io/5ekqr/?view_only=4416037a1b164e6287d95e7f24dd0a0a

