## Supplementary Information for "An ALE meta-analytic review of musical expertise"

+Shared first-authorship

*Corresponding author

**Search strings.**

Last search: March, 2021

**PubMed**

("music"[MeSH Terms] OR "musician"[All Fields] OR "music"[All Fields]) AND ("education"[Subheading] OR "education"[All Fields] OR "training"[All Fields] OR "education"[MeSH Terms] OR ("motor skills"[MeSH Terms] OR ("motor"[All Fields] AND "skills"[All Fields]) OR "motor skills"[All Fields])) AND ("magnetic resonance imaging"[MeSH Terms] OR ("magnetic"[All Fields] AND "resonance"[All Fields] AND "imaging"[All Fields]) OR "magnetic resonance imaging"[All Fields] OR "plasticity"[All Fields] OR "functional connectivity"[All Fields] OR "resting-state"[All Fields] OR "structural"[All Fields] OR ("brain"[MeSH Terms] AND "activity"[All Fields]))

719 results

**PsycInfo**

(((Music OR musician) AND (education OR training)) AND magnetic resonance imaging)

155 results

**Scopus**

(TITLE-ABS-KEY ( music ) OR TITLE-ABS-KEY (musician)) AND (TITLE-ABS-KEY (education) OR TITLE-ABS-KEY (training)) AND TITLE-ABS-KEY (magnetic resonance imaging)

295 results

**Supplementary Figure 1. PRISMA flowchart for literature search process.**

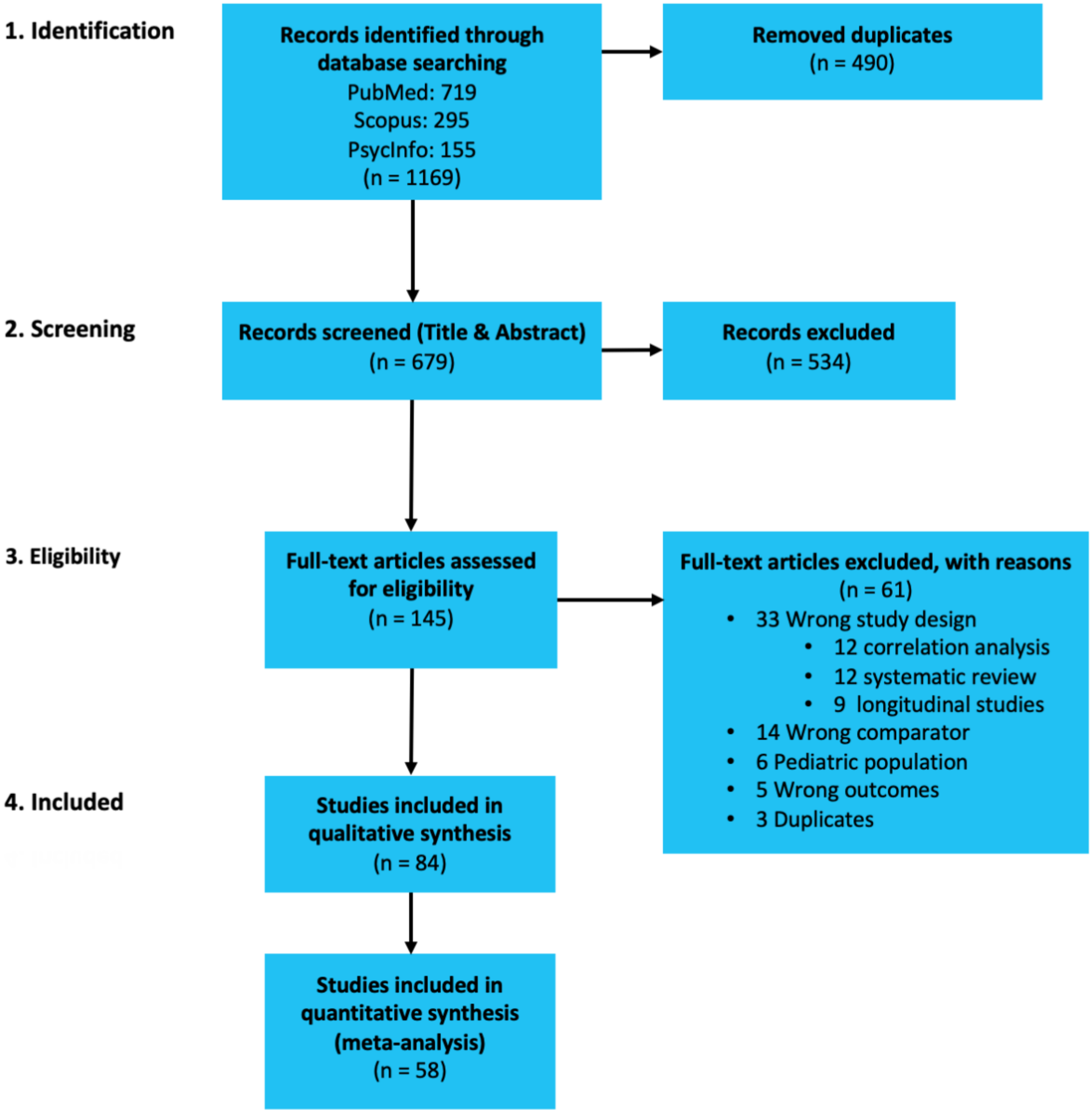

**Supplementary Figure 1**. PRISMA flowchart for literature search process.

**PRISMA Checklist**

SI = Supplementary Information

| Section/topic | # | Checklist item | Reported on page # |
|---|---|---|---|
| **TITLE** | | | |
| Title | 1 | Identify the report as a systematic review, meta-analysis, or both. | 1 |
| **ABSTRACT** | | | |
| Structured summary | 2 | Provide a structured summary including, as applicable: background; objectives; data sources; study eligibility criteria, participants, and interventions; study appraisal and synthesis methods; results; limitations; conclusions and implications of key findings; systematic review registration number. | 2 |
| **INTRODUCTION** | | | |
| Rationale | 3 | Describe the rationale for the review in the context of what is already known. | 3 |
| Objectives | 4 | Provide an explicit statement of questions being addressed with reference to participants, interventions, comparisons, outcomes, and study design (PICOS). | 3 |
| **METHODS** | | | |
| Protocol and registration | 5 | Indicate if a review protocol exists, if and where it can be accessed (e.g., Web address), and, if available, provide registration information including registration number. | 12 |
| Eligibility criteria | 6 | Specify study characteristics (e.g., PICOS, length of follow-up) and report characteristics (e.g., years considered, language, publication status) used as criteria for eligibility, giving rationale. | 12 |
| Information sources | 7 | Describe all information sources (e.g., databases with dates of coverage, contact with study authors to identify additional studies) in the search and date last searched. | 12 |
| Search | 8 | Present full electronic search strategy for at least one database, including any limits used, such that it could be repeated. | SI |
| Study selection | 9 | State the process for selecting studies (i.e., screening, eligibility, included in systematic review, and, if applicable, included in the meta-analysis). | 12 |
| Data collection process | 10 | Describe method of data extraction from reports (e.g., piloted forms, independently, in duplicate) and any processes for obtaining and confirming data from investigators. | 12 |
| Data items | 11 | List and define all variables for which data were sought (e.g., PICOS, funding sources) and any assumptions and simplifications made. | 12 |
| Risk of bias in individual studies | 12 | Describe methods used for assessing risk of bias of individual studies (including specification of whether this was done at the study or outcome level), and how this information is to be used in any data synthesis. | 12 |
| Summary measures | 13 | State the principal summary measures (e.g., risk ratio, difference in means). | 12 |
| Synthesis of results | 14 | Describe the methods of handling data and combining results of studies, if done, including measures of consistency (e.g., $I^2$) for each meta-analysis. | 12 |

| Section/topic | # | Checklist item | Reported on page # |
|---|---|---|---|
| Risk of bias across studies | 15 | Specify any assessment of risk of bias that may affect the cumulative evidence (e.g., publication bias, selective reporting within studies). | 12 |
| Additional analyses | 16 | Describe methods of additional analyses (e.g., sensitivity or subgroup analyses, meta-regression), if done, indicating which were pre-specified. | 12 |
| **RESULTS** | | | |
| Study selection | 17 | Give numbers of studies screened, assessed for eligibility, and included in the review, with reasons for exclusions at each stage, ideally with a flow diagram. | 4 |
| Study characteristics | 18 | For each study, present characteristics for which data were extracted (e.g., study size, PICOS, follow-up period) and provide the citations. | 4 |
| Risk of bias within studies | 19 | Present data on risk of bias of each study and, if available, any outcome level assessment (see item 12). | 4 |
| Results of individual studies | 20 | For all outcomes considered (benefits or harms), present, for each study: (a) simple summary data for each intervention group (b) effect estimates and confidence intervals, ideally with a forest plot. | 4 |
| Synthesis of results | 21 | Present results of each meta-analysis done, including confidence intervals and measures of consistency. | 4-6 |
| Risk of bias across studies | 22 | Present results of any assessment of risk of bias across studies (see Item 15). | 4 |
| Additional analysis | 23 | Give results of additional analyses, if done (e.g., sensitivity or subgroup analyses, meta-regression [see Item 16]). | 5 |
| **DISCUSSION** | | | |
| Summary of evidence | 24 | Summarize the main findings including the strength of evidence for each main outcome; consider their relevance to key groups (e.g., healthcare providers, users, and policy makers). | 7 |
| Limitations | 25 | Discuss limitations at study and outcome level (e.g., risk of bias), and at review-level (e.g., incomplete retrieval of identified research, reporting bias). | 9 |
| Conclusions | 26 | Provide a general interpretation of the results in the context of other evidence, and implications for future research. | 11 |
| **FUNDING** | | | |
| Funding | 27 | Describe sources of funding for the systematic review and other support (e.g., supply of data); role of funders for the systematic review. | 19 |

**Supplementary Table 1. Characteristics of MRI acquisition**

| | | | Teslas | MRI-system | MRI-model | Head-coil | T1 sequence | TR (ms) | TE (ms) | Voxel size (mm) | T2*sequence | TR (ms) | TE (ms) | Voxel size (mm) | DTI sequence | TR (ms) | TE (ms) | Voxel size (mm) |
|---|---|---|---|---|---|---|---|---|---|---|---|---|---|---|---|---|---|---|
| 1 | Abdul-K | 2011a | 1.5 | GE | Signa | quadrature | SPGR | 34 | 9 | - | - | - | - | - | - | - | - | - |
| 2 | Abdul-K | 2011b | 1.5 | Siemens | Symphony | 8-channel | MPRAGE | 1660 | 3.04 | 1x1x1 | - | - | - | - | DWI | 10100 | 106 | 2x2x2 |
| 3 | Acer | 2018 | 1.5 | Siemens | Aera | - | MPRAGE | 1900 | 2.84 | 1x1x1 | - | - | - | - | EPI | 3500 | 83 | 1.8x1.8x3.5 |
| 4 | Amunts | 1997 | 1.5 | Siemens | - | - | T1w | - | - | 1x1x1.17 | - | - | - | - | - | - | - | - |
| 5 | Angulo-P | 2014 | 3 | GE | Discovery | - | T1w | 2300 | 3 | 1x1x1 | EPI | 3000 | 40 | 2x2x3 | - | - | - | - |
| 6 | Bailey | 2014 | 3 | Siemens | Trio | 32-channel | T1w | 2300 | 2.98 | 1x1x1 | - | - | - | - | - | - | - | - |
| 7 | Bangert | 2006 | 1.5 | GE | Signa | quadrature | SPGR | - | - | - | EPI | 4500 | 40 | - | - | - | - | - |
| 8 | Baumann | 2007 | 3 | Philips | Intera | 8-channel | SPGR | 20 | 2.3 | 0.98x0.98x0.75 | EPI | 2000 | 35 | 2.75x2.75x2.75 | - | - | - | - |
| 9 | Bengtsson | 2005 | 1.5 | GE | Signa | - | T1w | 24 | 6 | 0.86x0.86x2 | - | - | - | - | EPI | 18000 | 107 | 1.8x1.8x3 |
| 10 | Berkowitz | 2010 | 3 | Philips | Intera | 8-channel | MPRAGE | - | - | 1x0.94x0.94 | EPI | 2500 | 35 | - | - | - | - | - |
| 11 | Bermudez | 2005 | - | - | - | - | T1w | - | - | - | - | - | - | - | - | - | - | - |
| 12 | Bermudez | 2009 | 1.5 | Siemens | Vision | - | T1w | 22 | 9.2 | 1x1x1 | - | - | - | - | - | - | - | - |
| 13 | Bianchi | 2017 | 3 | Philips | Achieva | 32-channel | T1w | 6056 | 2.78 | 0.85 | EPI | 10000 | 30 | 3x3x3 | - | - | - | - |
| 14 | Chen | 2008 | 1.5 | Siemens | Sonata | - | T1w | - | - | 1x1x1 | EPI | 10000 | 50 | 5x5x5 | - | - | - | - |
| 15 | Choi | 2015 | - | - | - | - | T1w | 1900 | 2.52 | 1x1x1 | - | - | - | - | - | - | - | - |
| 16 | De Manzano | 2018 | 3 | GE | Discovery | 8-channel | IRPFSGE | 6.7 | 2.9 | 1x1x1 | - | - | - | - | - | - | - | - |
| 17 | Du | 2017 | 3 | Siemens | Magnetom | 32-channel | MPRAGE | 2300 | 2.98 | 1x1x1 | EPI | 636 | 30 | 3x3x3 | - | - | - | - |
| 18 | Elmer | 2012 | 3 | Philips | Intera | 8-channel | - | - | - | - | EPI | 3000 | 35 | 1.72x1.72x4 | - | - | - | - |
| 19 | Elmer | 2013 | 3 | Philips | Achieva | 8-channel | T1w | 8.06 | 3.7 | 0.94x0.94x0.94 | - | - | - | - | - | - | - | - |
| 20 | Elmer | 2016 | 3 | Philips | Achieva | 8-channel | - | - | - | - | - | - | - | - | EPI | 13007 | 55 | 2x2x2 |
| 21 | Gaab | 2003 | 1.5 | Siemens | Vision | - | T1w | - | - | 1x1x1 | custom | 17000 | - | - | - | - | - | - |
| 22 | Gaab | 2006 | 3 | GE | Signa | - | - | - | - | - | EPI | - | - | - | - | - | - | - |
| 23 | Gagnepain | 2017 | 3 | Philips | Achieva | - | T1w | 20 | 4.6 | 1x1x1 | EPI | 500 | 80 | 2x1x1 | - | - | - | - |
| 25 | Gaser | 2003 | 1.5 | Siemens | Vision | - | MPRAGE | - | - | 1x1x1 | - | - | - | - | - | - | - | - |
| 24 | Gärtner | 2013 | 3 | Siemens | Magnetom | 12-channel | MPRAGE | 2250 | 3.03 | 1x1x1 | - | - | - | - | - | - | - | - |
| 26 | Giacosa | 2016 | 3 | Siemens | Trio | 32-channel | - | 9340 | 88 | 2x2x2 | - | - | - | - | - | - | - | - |
| 27 | Groussard | 2010 | 3 | Philips | Achieva | - | FFE | 20 | 4.6 | 1x1x1 | EPI | 2382 | 30 | 2.8x2.8x2.8 | - | - | - | - |
| 28 | Groussard | 2014 | 3 | Philips | Achieva | - | FFE | 20 | 4.6 | 1x1x1 | - | - | - | - | - | - | - | - |
| 29 | Halwani | 2011 | 3 | GE | - | - | T1w | - | - | 0.93x0.93x0.93 | - | - | - | - | EPI | 10000 | 86.9 | 2.5x2.5x2.6 |
| 30 | Han | 2009 | 3 | GE | - | 8-channel | SPGR | 8.5 | 3.4 | 1x1x1 | - | - | - | - | DWI | 10000 | 70.8 | - |
| 31 | Harris | 2015 | 3 | Philips | Intera | 8-channel | T1w | - | - | - | EPI | 16000 | 30 | 3.5x3.5x3.5 | - | - | - | - |
| 32 | Haslinger | 2004 | 1.5 | Philips | Intera | birdcage | T1w | - | - | - | EPI | 3000 | 50 | 3.59x3.59.5 | - | - | - | - |
| 33 | Haslinger | 2005 | 1.5 | Philips | Intera | birdcage | T1w | - | - | - | EPI | 3000 | 50 | 3.59x3.59.6 | - | - | - | - |
| 34 | Herdener | 2010 | 1.5 | Siemens | - | - | MPRAGE | - | - | 1.2x1x1 | EPI | 1850 | 61 | 5x5x5 | - | - | - | - |
| 35 | Herdener | 2014 | - | - | - | - | - | - | - | - | EPI | - | - | - | - | - | - | - |
| 36 | Hernández | 2019 | 3 | Philips | Achieva | - | MPRAGE | 8.4 | 3.8 | 0.9x0.89x0.8 | - | - | - | - | - | - | - | - |
| 37 | Huang | 2010 | 1.5 | Siemens | Sonata | custom | FLASH | 30 | 1.17 | - | EPI | 2000 | 60 | - | - | - | - | - |
| 38 | Hutchinson | 2003 | 1.5 | Siemens | Vision | - | T1w | - | - | 1x1x1 | - | - | - | - | - | - | - | - |
| 39 | Imfeld | 2009 | 3 | GE | Signa | 8-channel | - | - | - | - | - | - | - | - | EPI | 8000 | 91 | - |
| 40 | James | 2014 | 3 | Siemens | Trio | - | MPRAGE | 1900 | 2.27 | 1x1x1 | - | - | - | - | - | - | - | - |
| 41 | Karpati | 2017 | 3 | Siemens | Trio | 32-channel | T1w | 2300 | 2.98 | 1x1x1 | - | - | - | - | - | - | - | - |
| 42 | Kleber | 2010 | 1.5 | Siemens | Vision | - | MPRAGE | - | - | - | EPI | 10000 | 40 | 3 | - | - | - | - |
| 43 | Kleber | 2016 | 1.5 | Siemens | Sonata | 8-channel | MPRAGE | 1300 | 3.19 | - | - | - | - | - | - | - | - | - |
| 44 | Koelsch | 2005 | 3 | GE | - | - | T1w | - | - | 1x1x1.5 | - | 6000 | - | - | - | - | - | - |
| 45 | Koeneke | 2004 | 1.5 | GE | - | - | - | - | - | - | EPI | 2000 | 40 | 3.125x3.126x6 | - | - | - | - |
| 46 | Krings | 2000 | 1.5 | Philips | Gyroscan | - | T1w | - | - | - | EPI | 456 | 35 | - | - | - | - | - |
| 47 | Krishnan | 2018 | 1.5 | Siemens | Avanto | 32-channel | T1w | 2730 | 3.57 | 1x1x1 | EPI | 9500 | 50 | 2mm | - | - | - | - |
| 48 | Lee | 2011 | 3 | Siemens | Trio | - | T1w | 2300 | 9.38 | 1x1x1 | EPI | 3000 | 40 | 3x3x3.3 | - | - | - | - |

| | | | | | | | | | | | | | | | | | | |
|---|---|---|---|---|---|---|---|---|---|---|---|---|---|---|---|---|---|---|
| 49 | Limb | 2006 | 3 | GE | Signa | quadrature | T1w | - | - | - | EPI | 2000 | 30 | 6mm | - | - | - | - |
| 50 | Liu | 2018 | 3 | Siemens | Magnetom | - | T1w | 1900 | 2.52 | 1x1x1 | EPI | 2000 | 30 | 3x3.4x3.4 | - | - | - | - |
| 51 | Matsui | 2013 | 1.5 | GE | Signa | - | T1w | - | - | 1x1x1.5 | EPI | 3000 | 55 | 4mm | - | - | - | - |
| 52 | Mathews | 2020 | 3 | Siemens | Trio | 32-channel | T1w | 2420 | 3.7 | 1x1x1 | EPI | 2000 | 27.92 | 2.35x2.35x2.5 | - | - | - | - |
| 53 | Meister | 2005 | 1.5 | Philips | Gyroscan | quadrature | - | - | - | - | EPI | 3587 | 50 | 5mm | - | - | - | - |
| 54 | Morrison | 2003 | 1.5 | GE | - | - | T1w | - | - | - | EPI | 2500 | 50 | 3.5x3.5x5 | - | - | - | - |
| 55 | Oechslin | 2010 | 3 | GE | Signa | 8-channel | - | - | - | - | - | - | - | - | DWI | 8000 | 91 | - |
| 56 | Oechslin | 2013 | 3 | Siemens | Trio | - | MPRAGE | 1900 | 2.27 | 1x1x1 | EPI | 18600 | 30 | 3.2x3.2x3.2 | - | - | - | - |
| 57 | Oechslin | 2018 | 3 | Siemens | Trio | - | - | - | - | - | - | - | - | - | DWI | 8300 | 83 | 2x2x2 |
| 58 | Ohnishi | 2001 | 1.5 | Siemens | Magnetom | quadrature | - | - | - | - | EPI | 3000 | 60 | 3.44x3.44x4 | - | - | - | - |
| 59 | Ono | 2015 | 3 | Bruker | Medspec | birdcage | T1w | - | - | 1x1x1 | EPI | 2000 | 30 | 3mm | - | - | - | - |
| 60 | Öztürk | 2002 | 1.5 | GE | Signa | - | T1w | 500 | 20 | - | - | - | - | - | - | - | - | - |
| 61 | Park | 2014 | 3 | Siemens | Magnetom | TIM | MPRAGE | 2400 | 3.06 | 1x1x1 | EPI | 3000 | 30 | 3x3x4 | - | - | - | - |
| 62 | Pau | 2013 | 3 | Siemens | Magnetom | 12-channel | MPRAGE | - | - | 1x1x1 | EPI | 2000 | 30 | 3x3x3 | - | - | - | - |
| 63 | Petrini | 2011 | 3 | GE | Horizon | - | SPGR | - | - | 1.5x0.9x0.9 | EPI | 2000 | 35 | - | - | - | - | - |
| 64 | Rüber | 2015 | 3 | GE | - | - | T1w | - | - | 0.93x0.93x1.5 | - | - | - | - | - | - | - | - |
| 65 | Sakreida | 2018 | 3 | Siemens | Magnetom | 8-channel | T1w | 2040 | 5.57 | 1x1x1 | EPI | 2000 | 30 | 3x3x4.2 | - | - | - | - |
| 66 | Sato | 2015 | 3 | Philips | Achieva | - | MPRAGE | 15 | 3.3 | 1x1x1 | - | - | - | - | - | - | - | - |
| 67 | Schlaffke | 2020 | 3 | Philips | Achieva | 32-channel | MPRAGE | 8.3 | 3.8 | 1x1x1 | EPI | 2500 | 35 | 2x2x3 | - | - | - | - |
| 68 | Schlaug | 1995a | 1.5 | - | - | - | - | - | - | - | - | - | - | - | - | - | - | - |
| 69 | Schlaug | 1995b | 1.5 | - | - | - | - | - | - | - | - | - | - | - | - | - | - | - |
| 70 | Schlaug | 2005 | - | - | - | - | - | - | - | - | - | - | - | - | - | - | - | - |
| 71 | Schmithorst | 2002 | 3 | Bruker | Medspec | - | T1w | - | - | - | - | - | - | - | EPI | 6000 | 87 | - |
| 72 | Schmithorst | 2003 | 3 | Bruker | Biospec | - | T1w | - | - | - | EPI | 3000 | 38 | 5mm | - | - | - | - |
| 73 | Schmithorst | 2004 | 3 | Bruker | Medspec | - | T1w | - | - | - | EPI | 3000 | 38 | 5mm | - | - | - | - |
| 75 | Schneider | 2002 | 1.5 | Philips | Edge | - | T1w | - | - | 1x1x1 | - | - | - | - | - | - | - | - |
| 74 | Seung | 2005 | 1.5 | GE | Signa | - | T1w | - | - | - | EPI | 3000 | 60 | 3.75x3.75x5 | - | - | - | - |
| 76 | Sluming | 2002 | 1.5 | GE | Signa | quadrature | SPGR | 34 | 9 | - | - | - | - | - | - | - | - | - |
| 77 | Sluming | 2007 | 1.5 | GE | - | - | - | - | - | - | EPI | 3000 | 40 | 5mm | - | - | - | - |
| 78 | Steele | 2013 | 3 | Siemens | Trio | 32-channel | MPRAGE | 2300 | 2.98 | 1x1x1 | - | - | - | - | DWI | 9340 | 88 | 2x2x2 |
| 79 | Vaquero | 2016 | 3 | Siemens | Magnetom | - | MPRAGE | 16 | 4.9 | 1x1x1 | - | - | - | - | - | - | - | - |
| 80 | Vaquero | 2020 | 3 | Siemens | Magnetom | - | T1w | - | - | - | - | - | - | - | EPI | 1000 | 90 | 2x2x2 |
| 81 | Wang | 2019 | 3 | Siemens | Magnetom | - | MPRAGE | 2300 | 3.24 | 1x1x1 | - | - | - | - | - | - | - | - |
| 82 | Zarate | 2005 | 1.5 | - | - | - | T1w | - | - | - | - | 10000 | 2.125 | - | - | - | - | - |
| 83 | Zarate | 2008 | 1.5 | Siemens | Sonata | - | T1w | - | - | 1x1x1 | EPI | 10000 | 85 | 5x5x5 | - | - | - | - |
| 84 | Zuk | 2014 | 3 | Siemens | Trio | - | - | - | - | - | EPI | 2000 | 30 | 3x3x4 | - | - | - | - |

GM, grey matter; WM, white matter; MRI, magnetic resonance imaging; FFE, fast field echo sequence; FLASH, fast low angle shot sequence; FSL, functional MRI of the brain software library; GE, gradient echo pulse; IR-FSPGR, fast spoiled gradient sequence with inversion preparation; MPRAGE, magnetization-prepared rapid acquisition with gradient echo sequence; MDEFT, modified driven equilibrium Fourier transform; SPGR, spoiled gradient recalled sequence; SPM, statistical parametric mapping; TFE, turbo field echo sequence; VBM, voxel-based morphometry[1,2].

**Supplementary Table 2. Characteristics of MRI analyses.**

| | | | VBM Analysis Software | VBM Analysis Method | DTI Analysis Software | DTI Analysis Method | CT Analysis Software | CT Analysis Method | fMRI Analysis Method | fMRI Analysis software | Task | Stimuli | Control |
|---|---|---|---|---|---|---|---|---|---|---|---|---|---|
| 1 | Abdul-K | 2011a | SPSS 16 | VBM-ROI | - | - | - | - | - | - | - | - | - |
| 2 | Abdul-K | 2011b | - | - | SPSS | FA | - | - | - | - | - | - | - |
| 3 | Acer | 2018 | VBM8 | VBM-whole brain | DTIStudio | FA+MD | - | - | - | - | - | - | - |
| 4 | Amunts | 1997 | - | VBM-ROI | - | - | - | - | - | - | - | - | - |
| 5 | Angulo-P | 2014 | - | - | - | - | - | - | GLM | FSL | listen | music excerpts | speech |
| 6 | Bailey | 2014 | FSL | VBM-whole brain | - | - | FSL | CT | - | - | - | - | - |
| 7 | Bangert | 2006 | - | - | - | - | - | - | GLM | SPM99 | listen+play | piano | rest |
| 8 | Baumann | 2007 | - | - | - | - | - | - | GLM | SPM2 | listen+tapping | music excerpts | rest |
| 9 | Bengtsson | 2005 | SPM99 | VBM-whole brain | SPM-99 | FA | - | - | - | - | - | - | - |
| 10 | Berkowitz | 2010 | - | - | - | - | - | - | GLM | BrainVoyager | play music | piano | patterns |
| 11 | Bermudez | 2005 | - | VBM-whole brain | - | - | - | - | - | - | - | - | - |
| 12 | Bermudez | 2009 | CIVET | VBM-whole brain | - | - | CIVET | MACACC | - | - | - | - | - |
| 13 | Bianchi | 2017 | - | - | - | - | - | - | GLM | SPM8 | listen+rate | tones | rest |
| 14 | Chen | 2008 | - | - | - | - | - | - | GLM | fMRISTAT | listen+tapping | rhythm | rest |
| 15 | Choi | 2015 | Brainvoyager | - | - | - | BrainVoyager | CT | - | - | - | - | - |
| 16 | De Manzano | 2018 | SPM12 | VBM-ROI | Mrtrix3 | FA | FreeSurfer | LME | - | - | - | - | - |
| 17 | Du | 2017 | - | - | - | - | - | - | GLM | AFNI | listen+rate | speech | noise |
| 18 | Elmer | 2012 | - | - | - | - | - | - | GLM | SPM8 | listen+rate | speech | noise |
| 19 | Elmer | 2013 | - | - | - | - | FreeSurfer | - | - | - | - | - | - |
| 20 | Elmer | 2016 | - | - | FSL | FA | - | - | - | - | - | - | - |
| 21 | Gaab | 2003 | - | - | - | - | - | - | GLM | SPM99 | listen+rate | tones | motor |
| 22 | Gaab | 2006 | - | - | - | - | - | - | GLM | SPM2 | listen+play | tones | - |
| 23 | Gagnepain | 2017 | - | - | - | - | - | - | GLM | SPM8 | listen+rate | melodies | proverbs |
| 25 | Gaser | 2003 | SPM99 | VBM-whole brain | - | - | - | - | - | - | - | - | - |
| 24 | Gärtner | 2013 | SPM8 | DBM-ROI | - | - | - | - | - | - | - | - | - |
| 26 | Giacosa | 2016 | - | - | FSL | TBSS | - | - | - | - | - | - | - |
| 27 | Groussard | 2010 | SPM5 | VBM-whole brain | - | - | - | - | GLM | SPM5 | - | melodies | unfamiliar melodies |
| 28 | Groussard | 2014 | SPM12 | VBM-whole brain | - | - | - | - | - | - | - | - | - |
| 29 | Halwani | 2011 | - | - | FSL | FA | - | - | - | - | - | - | - |
| 30 | Han | 2009 | SPM2 | VBM-whole brain | AFNI | FA | - | - | - | - | - | - | - |
| 31 | Harris | 2015 | - | - | - | - | - | - | GLM | SPM5 | listen+imagine | piano | score-reading |
| 32 | Haslinger | 2004 | - | - | - | - | - | - | GLM | SPM99 | play music | piano | rest |
| 33 | Haslinger | 2005 | - | - | - | - | - | - | GLM | SPM99 | audiovisual | piano | rest |
| 34 | Herdener | 2010 | - | - | - | - | - | - | GLM | BrainVoyager | - | tones | mismatch |
| 35 | Herdener | 2014 | - | - | - | - | - | - | GLM | BrainVoyager | listen+imagine | tones | mismatch |
| 36 | Hernández | 2019 | SPM12 | VBM-ROI | - | - | - | - | - | - | - | - | - |
| 37 | Huang | 2010 | - | - | - | - | - | - | GLM | AFNI | listen+rate | words | - |
| 38 | Hutchinson | 2003 | custom | VBM-ROI | - | - | - | - | - | - | - | - | - |
| 39 | Imfeld | 2009 | - | - | SPM5 | FA | - | - | - | - | - | - | - |
| 40 | James | 2014 | SPM8 | VBM-whole brain | - | - | - | - | - | - | - | - | - |
| 41 | Karpati | 2017 | CIVET | VBM-whole brain | - | - | - | - | - | - | - | - | - |
| 42 | Kleber | 2010 | - | - | - | - | - | - | GLM | SPM5 | singing | - | rest |
| 43 | Kleber | 2016 | SPM5 | VBM-whole brain | - | - | - | - | - | - | - | - | - |
| 44 | Koelsch | 2005 | - | - | - | - | - | - | GLM | SPM99 | listen | tones | deviants |
| 45 | Koeneke | 2004 | - | - | - | - | - | - | GLM | SPM99 | play | - | rest |
| 46 | Krings | 2000 | - | - | - | - | - | - | GLM | - | tapping | - | rest |

| | | | | | | | | | | | | | |
|---|---|---|---|---|---|---|---|---|---|---|---|---|---|
| 47 | Krishnan | 2018 | FSL | VBM-whole brain | - | - | - | - | GLM | SPM8 | listen | beatbox/guitar | rest |
| 48 | Lee | 2011 | - | - | - | - | - | - | GLM | SPM8 | audiovisual | sentences/piano | rest |
| 49 | Limb | 2006 | - | - | - | - | - | - | GLM | SPM99 | listen | rhythm | rest |
| 50 | Liu | 2018 | - | - | - | - | - | - | GLM | SPM8 | listen+rate | music excerpts | rest |
| 51 | Matsui | 2013 | - | - | - | - | - | - | GLM | SPM8 | listen | music excerpts | scrambled |
| 52 | Mathews | 2020 | - | - | - | - | - | - | GLM | SPM12 | listen+rate | piano | rest |
| 53 | Meister | 2005 | - | - | - | - | - | - | GLM | SPM99 | play music | - | rest |
| 54 | Morrison | 2003 | - | - | - | - | - | - | GLM | MEDx | listen | music excerpts | speech |
| 55 | Oechslin | 2010 | - | - | SPM5 | FA+MD | - | - | - | - | - | - | - |
| 56 | Oechslin | 2013 | - | - | - | - | - | - | GLM | SPM8 | listen+rate | music excerpts | scrambled |
| 57 | Oechslin | 2018 | - | - | FSL | FA | - | - | - | - | - | - | - |
| 58 | Ohnishi | 2001 | - | - | - | - | - | - | GLM | SPM99 | listen | music excerpts | rest |
| 59 | Ono | 2015 | - | - | - | - | - | - | GLM | SPM8 | tapping | conductors | metronome |
| 60 | Öztürk | 2002 | - | - | - | - | - | - | - | - | - | - | - |
| 61 | Park | 2014 | - | - | - | - | - | - | GLM | SPM8 | listen+rate | music excerpts | noise |
| 62 | Pau | 2013 | - | - | - | - | - | - | GLM | SPM5 | tapping | tones | noise |
| 63 | Petrini | 2011 | - | - | - | - | - | - | GLM | BrainVoyager | listen+rate | drums | rest |
| 64 | Rüber | 2015 | - | - | FSL | FA | - | - | - | - | - | - | - |
| 65 | Sakreida | 2018 | - | - | - | - | - | - | GLM | SPM8 | tapping | rhythm | rest |
| 66 | Sato | 2015 | SPM12 | VBM-whole brain | - | - | - | - | - | - | - | - | - |
| 67 | Schlaffke | 2020 | - | - | ExploreDTI | FA+MD | - | - | GLM | SPM8 | tapping | drums | rest |
| 68 | Schlaug | 1995a | - | - | - | - | - | - | - | - | - | - | - |
| 69 | Schlaug | 1995b | - | - | - | - | - | - | - | - | - | - | - |
| 70 | Schlaug | 2005 | - | - | - | - | - | - | - | - | - | - | - |
| 71 | Schmithorst | 2002 | - | - | IDL | FA | - | - | - | - | - | - | - |
| 72 | Schmithorst | 2003 | - | - | - | - | - | - | GLM | IDL | listen | melodies | rest |
| 73 | Schmithorst | 2004 | - | - | - | - | - | - | GLM | IDL | math | visual | rest |
| 75 | Schneider | 2002 | Brainvoyager | VBM-ROI | - | - | - | - | - | - | - | - | - |
| 74 | Seung | 2005 | - | - | - | - | - | - | GLM | SPM99 | listen | music excerpts | rest |
| 76 | Sluming | 2002 | SPM99 | VBM-whole brain | - | - | - | - | - | - | - | - | - |
| 77 | Sluming | 2007 | - | - | - | - | - | - | GLM | SPM99 | visual | drawing | rest |
| 78 | Steele | 2013 | - | - | FSL | FA | - | - | - | - | - | - | - |
| 79 | Vaquero | 2016 | SPM8 | VBM-whole brain | - | - | - | - | - | - | - | - | - |
| 80 | Vaquero | 2020 | - | - | FSL | FA+RD | - | - | - | - | - | - | - |
| 81 | Wang | 2019 | SPM12 | VBM-whole brain | - | - | - | - | - | - | - | - | - |
| 82 | Zarate | 2005 | - | - | - | - | - | - | - | - | sing | - | - |
| 83 | Zarate | 2008 | - | - | - | - | - | - | GLM | fMRISTAT | sing | vocal waves | noise |
| 84 | Zuk | 2014 | - | - | - | - | - | - | - | - | listen+rate | sounds | - |

fMRI, functional magnetic resonance imaging; VBM, voxel-based morphometry; DTI, diffusion tensor imaging; CT, cortical thickness; ROI, region-of-interest; FSL, functional MRI of the brain software library; MPRAGE, magnetization-prepared rapid acquisition with gradient echo sequence; SPGR, spoiled gradient recalled sequence; SPM, statistical parametric mapping; FA, fractional anisotropy.

**Supplementary Table 3.** **Summary of MRI quality.**

| | | | MRI design described | Age reported | Sample gender reported | Sample handedness reported | Matched control group | Ethics approval reported | Image acquisition described | Image processing described | Statistical MRI-analysis described | Software package specified | Multiple comparison correction described | Figures and tables | Main outcomes: M > NM | NM > M |
|---|---|---|---|---|---|---|---|---|---|---|---|---|---|---|---|---|
| 1 | Abdul-K | 2011a | Y | Y | Y | Y | a,b,c | Y | Y | Y | Y | Y | Y | Y | lIFG | - |
| 2 | Abdul-K | 2011b | Y | Y | Y | Y | a,b,c | Y | Y | Y | Y | Y | Y | Y | rSCP, rMCP, rCRBL | - |
| 3 | Acer | 2018 | Y | Y | Y | Y | a,b | Y | Y | Y | Y | Y | N | Y | rCST, rCRBL, lCRBL, SMG, AnG, lSPL, lIPL, MidTG | - |
| 4 | Amunts | 1997 | Y | Y | Y | Y | a,b,c | N | Y | U | U | U | U | Y | PreCG | - |
| 5 | Angulo-P | 2014 | Y | Y | Y | Y | a,b,c | Y | Y | Y | Y | Y | Y | Y | PT | - |
| 6 | Bailey | 2014 | Y | Y | N | Y | c | Y | Y | Y | Y | Y | Y | Y | PMC, SI | - |
| 7 | Bangert | 2006 | Y | Y | Y | Y | a,b,c | N | Y | Y | Y | Y | Y | Y | MidTG, SFG, SRG, PreCG, IFG, IPL, HIPP, DLPFC, SMG, ACC, | - |
| 8 | Baumann | 2007 | Y | Y | Y | Y | b,c,e | Y | U | Y | Y | Y | N | Y | PMC, SMA | - |
| 9 | Bengtsson | 2005 | Y | Y | Y | Y | a,b,c | Y | Y | U | Y | Y | Y | Y | IC | - |
| 10 | Berkowitz | 2010 | Y | U | U | N | a | N | Y | Y | Y | Y | Y | Y | rTPJ | - |
| 11 | Bermudez | 2005 | N | N | N | N | - | N | U | N | U | N | Y | Y | STG, PT | - |
| 12 | Bermudez | 2009 | Y | Y | Y | Y | a,b,c | Y | Y | Y | Y | N | Y | Y | HG, PreCG, IFG | - |
| 13 | Bianchi | 2017 | Y | Y | Y | Y | a,b,c | Y | Y | Y | Y | Y | Y | Y | STG, HG, PP, IFG, PreCG, FusG, CRBL, IF, INS | - |
| 14 | Chen | 2008 | Y | Y | N | Y | c | Y | Y | Y | Y | Y | N | Y | DLPFC, IFG, CRBL | - |
| 15 | Choi | 2015 | Y | Y | Y | N | a,b | Y | U | Y | Y | Y | Y | Y | SI (lips) | SI (tongue) |
| 16 | De Manzano | 2018 | Y | Y | Y | N | a,b | Y | Y | Y | Y | Y | Y | Y | STG, CRBL, WM | - |
| 17 | Du | 2017 | Y | Y | Y | Y | a,b,c | Y | Y | Y | Y | Y | Y | Y | AnG, MidTG, IFG, | CRBL |
| 18 | Elmer | 2012 | Y | Y | Y | Y | a,b,c | Y | Y | Y | Y | Y | Y | Y | PT, PMC | - |
| 19 | Elmer | 2013 | Y | Y | Y | Y | a,b,c,d | Y | Y | Y | Y | Y | Y | Y | PT | - |
| 20 | Elmer | 2016 | Y | Y | Y | Y | a,b,c,d | Y | Y | Y | Y | Y | Y | Y | CC | - |
| 21 | Gaab | 2003 | Y | Y | Y | Y | a,b,c | N | Y | Y | Y | Y | Y | Y | PT, SMG, SPL, SPL, IFG | PT, CRBL, HIPP |
| 22 | Gaab | 2006 | Y | Y | Y | Y | a,b | N | U | N | Y | Y | Y | Y | - | lIFG, MidFG, ACC, IPL |
| 23 | Gagnepain | 2017 | Y | Y | Y | Y | a,b,c | Y | Y | Y | Y | Y | Y | Y | HIPP | - |
| 25 | Gaser | 2003 | Y | Y | Y | Y | a,b,c,d | Y | Y | U | U | Y | Y | Y | ITG, M1, SPL, HG, IFG, MidFG, CRBL | - |
| 24 | Gärtner | 2013 | Y | Y | Y | Y | a,b,c | Y | Y | Y | Y | Y | N | Y | CC, CST, RN, EnC, THA | - |
| 26 | Giacosa | 2016 | Y | Y | Y | N | a,b | Y | Y | Y | Y | Y | Y | Y | CST, IFOF, SFL, ILF | - |
| 27 | Groussard | 2010 | Y | Y | Y | Y | a,b,c,d | Y | Y | Y | Y | Y | N | Y | HIPP, CalC, LG, OFC, MCC, STG, CRBL | - |
| 28 | Groussard | 2014 | Y | Y | Y | Y | a,b,c | Y | Y | Y | Y | Y | N | Y | HIPP, SMA, SFG, MidFG, PCC, INS, STG | - |
| 29 | Halwani | 2011 | Y | Y | N | N | a | Y | Y | Y | Y | Y | Y | Y | AF | - |
| 30 | Han | 2009 | Y | Y | Y | Y | a,b,c | Y | Y | Y | Y | Y | N | Y | M1, SI, CRBL, IFG, IC, MB | OFC, ACC |
| 31 | Harris | 2015 | Y | Y | Y | Y | a,b,c | Y | Y | Y | Y | Y | Y | Y | PMC, SMG, PCC, STG, STS | - |
| 32 | Haslinger | 2004 | Y | Y | Y | Y | a,b,c | Y | Y | Y | Y | Y | Y | Y | - | PMC, ACC, CRBL ITG, LG, CAU, SMA MidFG, PMC |
| 33 | Haslinger | 2005 | Y | Y | Y | Y | a,b,c | Y | Y | Y | Y | Y | Y | Y | PMC, SMA, IFG, MidFG, STG, M1, S1 | - |
| 34 | Herdener | 2010 | Y | Y | Y | N | a,b | Y | Y | Y | Y | Y | Y | Y | HIPP, PCN, INS | - |
| 35 | Herdener | 2014 | Y | Y | Y | N | a,b | Y | N | N | U | Y | Y | Y | SMG, INS | - |
| 36 | Hernández | 2019 | Y | Y | Y | N | a,b | Y | Y | Y | Y | Y | Y | Y | CAU | - |
| 37 | Huang | 2010 | Y | Y | Y | Y | a,b,c,d | Y | Y | Y | Y | Y | Y | Y | LG, IFG, HIPP, AMYG, MedFG. | - |
| 38 | Hutchinson | 2003 | Y | Y | Y | Y | a,b,c | Y | U | U | Y | Y | Y | Y | CRBL | - |
| 39 | Imfeld | 2009 | Y | Y | Y | Y | a,c | Y | Y | Y | Y | Y | N | Y | CST | - |
| 40 | James | 2014 | Y | Y | N | Y | a,c | Y | Y | Y | Y | Y | Y | Y | FusG, OFC, CRBL, IFG, IPL, HG | PostCG, PCN, CAU |
| 41 | Karpati | 2017 | Y | Y | Y | N | a,b | Y | Y | Y | Y | Y | Y | Y | STG, STS, MTG | - |
| 42 | Kleber | 2010 | Y | Y | Y | Y | c | Y | Y | Y | Y | Y | Y | Y | M1, SI, SMA, DLPFC, TP, GP, CRBL , PCN, PUT, THA | - |
| 43 | Kleber | 2016 | Y | Y | Y | Y | a,b,c | Y | Y | Y | Y | Y | Y | Y | SI, SMG, SII, STG, HIPP, CAU | - |
| 44 | Koelsch | 2005 | Y | Y | Y | Y | a,b,c | Y | Y | Y | Y | Y | N | Y | STG, SMG, PL, IFG | - |
| 45 | Koeneke | 2004 | Y | Y | Y | Y | a,b,c | Y | Y | Y | Y | Y | Y | Y | rCRBL | lCRBL, VER |
| 46 | Krings | 2000 | Y | Y | Y | Y | a,b,c | N | U | N | U | N | N | Y | - | S1, SMA, PMC, SPL |

| | | | | | | | | | | | | | | | | |
|---|---|---|---|---|---|---|---|---|---|---|---|---|---|---|---|---|
| 47 | Krishnan | 2018 | Y | Y | Y | N | a,b | Y | Y | Y | Y | Y | Y | Y | CRBL, IFG, IPC | IFG, ITG, PostCG, SMA, CRBL |
| 48 | Lee | 2011 | Y | Y | N | N | a | N | Y | Y | Y | Y | Y | Y | STS, MTG, PMC, CRBL | - |
| 49 | Limb | 2006 | Y | Y | Y | Y | a,b,c | Y | Y | Y | Y | Y | N | Y | MTG, AnG, SMG, FO, MidFG, SFG | STG, SMG, PCN, Cuneus, PreCG, GP, PUT |
| 50 | Liu | 2018 | Y | N | Y | Y | a,c | Y | Y | N | Y | Y | Y | Y | IPL, STG, HG, PCN, MTG, PCC | - |
| 51 | Matsui | 2013 | Y | Y | Y | Y | a,b,c | Y | Y | Y | Y | Y | Y | Y | MTG, STG | STG |
| 52 | Mathews | 2020 | Y | Y | Y | Y | a | Y | Y | Y | Y | Y | Y | Y | PreCG, SMA, IFG, HG, CAU, STG, SMG | - |
| 53 | Meister | 2005 | Y | Y | Y | Y | b,c | Y | Y | Y | Y | Y | U | Y | FP | - |
| 54 | Morrison | 2003 | Y | Y | Y | Y | b | Y | Y | Y | Y | Y | Y | Y | STG | - |
| 55 | Oechslin | 2010 | Y | Y | Y | Y | a,b | Y | Y | Y | Y | Y | Y | Y | AF | - |
| 56 | Oechslin | 2013 | Y | Y | N | Y | a,c | N | Y | Y | Y | Y | Y | Y | ACC, SMG, SMA, PO, PCN, PostCG | - |
| 57 | Oechslin | 2018 | Y | Y | Y | Y | a,b,c,d | N | Y | Y | Y | Y | Y | Y | rVentral stream | - |
| 58 | Ohnishi | 2001 | Y | Y | Y | Y | a,b,c | Y | Y | Y | Y | Y | Y | Y | STG, MidFG | - |
| 59 | Ono | 2015 | Y | Y | Y | Y | a,b,c | Y | Y | Y | Y | Y | Y | Y | SFG | - |
| 60 | Öztürk | 2002 | Y | Y | Y | Y | b,c | Y | U | U | Y | N | N | Y | CC | - |
| 61 | Park | 2014 | Y | Y | Y | Y | a,b,c | Y | Y | Y | Y | Y | N | Y | MidFG, PreCG, SFG, SMG, PostCG, IPL | - |
| 62 | Pau | 2013 | Y | Y | Y | Y | a,b,c | Y | Y | Y | Y | Y | Y | Y | SI, M1, SMA, PMC, SPL, IFG, CRBL, DLPFC, OL, INS, STG, MTG | S1, M1, SMA, PMC, IPL, IFG, CRBL, PUT, DLPFC, INS, STG, MTG |
| 63 | Petrini | 2011 | Y | Y | Y | Y | a,b,c | N | Y | Y | Y | Y | Y | Y | - | MidFG |
| 64 | Rüber | 2015 | Y | Y | Y | Y | a,b,c | Y | Y | Y | Y | Y | Y | Y | M1 | - |
| 65 | Sakreida | 2018 | Y | Y | Y | Y | a,b | Y | Y | Y | Y | Y | Y | Y | IPL, SMG, ITG,, MTG, MOG, IFG, MidFG, MidTG, SMA, SFG, PCN, SPL, PUT, INS, CRBL | IPL, AnG, SMG, CUN, IFG |
| 66 | Sato | 2015 | Y | Y | Y | Y | a,b,c | Y | Y | Y | Y | Y | N | Y | MOG, IFG, LG, STG, PCN, CAU, SPL, TP | CAU |
| 67 | Schlaffke | 2020 | Y | Y | Y | Y | a,b,c | Y | Y | Y | Y | Y | Y | Y | CC | - |
| 68 | Schlaug | 1995a | Y | Y | Y | Y | a,b | N | N | N | Y | N | Y | Y | CC | - |
| 69 | Schlaug | 1995b | Y | Y | Y | Y | a,b | N | N | N | Y | N | Y | Y | PT | - |
| 70 | Schlaug | 2005 | N | N | N | N | - | N | N | N | N | N | N | Y | SI, M1, PMC, SPL, HG, CRBL, IFG | - |
| 71 | Schmithorst | 2002 | Y | Y | N | N | a | Y | Y | Y | Y | Y | Y | Y | CC | - |
| 72 | Schmithorst | 2003 | Y | Y | Y | N | a | Y | Y | Y | Y | Y | Y | Y | FusG, LG | ACC, SFG |
| 73 | Schmithorst | 2004 | Y | Y | Y | N | a | Y | Y | Y | Y | Y | Y | Y | FusG, MedFG | IOG, MOG, ThA, OFG, IPL |
| 75 | Schneider | 2002 | Y | Y | Y | Y | a,b,c | Y | Y | Y | Y | Y | N | Y | HG | - |
| 74 | Seung | 2005 | Y | Y | Y | Y | a,b,c | Y | Y | Y | Y | Y | Y | Y | STG, MTG, IFG, SMG | PCN, LG |
| 76 | Sluming | 2002 | Y | Y | Y | Y | a,b,c | N | Y | Y | Y | Y | Y | Y | IFG | - |
| 77 | Sluming | 2007 | Y | Y | Y | Y | a,b,c | Y | Y | Y | Y | Y | Y | Y | IFG, AnG, ACC | PCN, AnG, SPL, PrecG, SMA |
| 78 | Steele | 2013 | Y | Y | Y | Y | a,c | Y | Y | Y | Y | Y | Y | Y | CC | - |
| 79 | Vaquero | 2016 | Y | Y | Y | Y | a,c | Y | Y | Y | Y | Y | Y | Y | PUT, AMYG, CS, LG, PUT, THA, STG | SMG, STG, PostCG |
| 80 | Vaquero | 2020 | Y | Y | Y | Y | a,b,c,d | Y | Y | Y | Y | Y | Y | Y | AF | - |
| 81 | Wang | 2019 | Y | Y | Y | Y | a,b,c,d | Y | Y | Y | Y | Y | Y | Y | - | INS |
| 82 | Zarate | 2005 | U | N | Y | N | a,b | N | Y | N | N | N | N | Y | ACC, STS, INS; PUT, SMA | ACC, IPL |
| 83 | Zarate | 2008 | Y | Y | Y | Y | a | Y | Y | Y | Y | Y | Y | Y | GP; CRBLV | ACC |
| 84 | Zuk | 2014 | Y | Y | Y | Y | a,b,c | Y | Y | Y | Y | Y | Y | Y | SMA, PFC | - |

Y, yes; N, no; U, unclear.

* a, age; b, sex; c, handedness; d, education.

MRI guidelines[1,2]

**Supplementary Table 4. Meta-analytic connectivity modelling of regions-of-interest resulted from structural and functional ALE meta-analyses.**

| Cluster Number | Volume (mm3) | MNI coordinates x | y | z | ALE | P | Z | Label (Side region BA) |
|---|---|---|---|---|---|---|---|---|
| **a. STRUCTURAL ALE ROIs** | | | | | | | | |
| *M>NM (GM)* | | | | | | | | |
| *1. STG-R BA13: 992 foci, 46 experiments, 593 subjects (x=50, y=-20, z=8)* | | | | | | | | |
| 1 | 24288 | -58 | -22 | 8 | 5E-02 | 2E-15 | 7.8 | L Superior Temporal Gyrus BA41 |
| | | -50 | -16 | 0 | 5E-02 | 7E-15 | 7.7 | L Superior Temporal Gyrus BA22 |
| | | -42 | -26 | 10 | 5E-02 | 7E-14 | 7.4 | L Transverse Temporal Gyrus BA41 |
| | | -38 | -32 | 14 | 5E-02 | 5E-13 | 7.1 | L Superior Temporal Gyrus BA41 |
| | | -50 | 10 | 2 | 4E-02 | 6E-10 | 6.1 | L Insula BA13 |
| | | -52 | -38 | 18 | 4E-02 | 1E-09 | 6.0 | L Insula BA13 |
| | | -32 | 20 | 4 | 4E-02 | 1E-09 | 6.0 | L Insula BA13 |
| | | -58 | -8 | -2 | 3E-02 | 6E-07 | 4.8 | L Superior Temporal Gyrus BA22 |
| | | -36 | 2 | 4 | 2E-02 | 2E-04 | 3.6 | L Claustrum |
| 2 | 14968 | 50 | -20 | 8 | 2E-01 | 0E+00 | 20.0 | R Superior Temporal Gyrus BA13 |
| | | 62 | -28 | 4 | 4E-02 | 4E-11 | 6.5 | R Superior Temporal Gyrus BA22 |
| | | 60 | -4 | -6 | 3E-02 | 6E-08 | 5.3 | R Superior Temporal Gyrus BA22 |
| | | 62 | -4 | 16 | 2E-02 | 2E-04 | 3.5 | R Precentral Gyrus BA4 |
| 3 | 5520 | 48 | 10 | 2 | 4E-02 | 7E-11 | 6.4 | R Precentral Gyrus BA44 |
| | | 38 | 22 | 0 | 3E-02 | 4E-08 | 5.4 | R Insula BA13 |
| | | 50 | 20 | -6 | 2E-02 | 2E-05 | 4.1 | R Inferior Frontal Gyrus BA47 |
| 4 | 3528 | -4 | 0 | 62 | 4E-02 | 4E-09 | 5.8 | L Medial Frontal Gyrus BA6 |
| | | 0 | 8 | 52 | 3E-02 | 5E-06 | 4.4 | L Medial Frontal Gyrus BA6 |
| | | 2 | 16 | 40 | 2E-02 | 9E-05 | 3.8 | L Cingulate Gyrus BA32 |
| 5 | 1856 | -30 | -56 | -28 | 3E-02 | 3E-07 | 5.0 | L Culmen |
| | | -14 | -62 | -18 | 3E-02 | 7E-06 | 4.3 | L Declive |
| | | -24 | -66 | -18 | 2E-02 | 1E-04 | 3.6 | L Declive |
| 6 | 1536 | -20 | 6 | 4 | 3E-02 | 7E-08 | 5.3 | L Lentiform Nucleus |
| | | -12 | -2 | 12 | 2E-02 | 5E-05 | 3.9 | L Caudate |
| 7 | 1472 | -12 | -18 | 2 | 3E-02 | 3E-08 | 5.4 | L Thalamus |
| *2. STG-L BA41: 1428 foci, 71 experiments, 961 subjects (x=-56, y=-20, z=2)* | | | | | | | | |
| 1 | 25008 | -56 | -20 | 2 | 3E-01 | 0E+00 | 25.6 | L Superior Temporal Gyrus BA41 |
| | | -56 | -2 | -8 | 5E-02 | 6E-12 | 6.8 | L Superior Temporal Gyrus BA22 |
| | | -40 | -34 | 14 | 3E-02 | 3E-07 | 5.0 | L Transverse Temporal Gyrus BA41 |
| | | -36 | 22 | 4 | 3E-02 | 2E-06 | 4.6 | L Insula BA13 |
| | | -46 | 22 | -10 | 3E-02 | 6E-06 | 4.4 | L Inferior Frontal Gyrus BA47 |
| | | -34 | 26 | -8 | 2E-02 | 7E-05 | 3.8 | L Insula BA13 |
| | | -56 | 6 | 10 | 2E-02 | 7E-05 | 3.8 | L Precentral Gyrus BA6 |
| | | -54 | 18 | 8 | 2E-02 | 2E-04 | 3.5 | L Inferior Frontal Gyrus BA44 |
| | | -58 | -4 | 20 | 2E-02 | 2E-04 | 3.5 | L Postcentral Gyrus BA4 |
| | | -58 | -8 | 26 | 2E-02 | 3E-04 | 3.4 | L Precentral Gyrus BA4 |
| | | -60 | -8 | 14 | 2E-02 | 4E-04 | 3.4 | L Precentral Gyrus BA43 |
| 2 | 19728 | 60 | -20 | 2 | 1E-01 | 5E-33 | 11.9 | R Superior Temporal Gyrus BA41 |
| | | 54 | 10 | -14 | 4E-02 | 3E-10 | 6.2 | R Superior Temporal Gyrus BA22 |
| 3 | 3416 | 36 | 22 | -6 | 4E-02 | 3E-09 | 5.8 | R Insula |
| | | 46 | 24 | 10 | 3E-02 | 5E-07 | 4.9 | R Inferior Frontal Gyrus BA45 |
| 4 | 2568 | -46 | 12 | 24 | 3E-02 | 7E-07 | 4.8 | L Inferior Frontal Gyrus BA9 |
| | | -54 | 22 | 20 | 3E-02 | 5E-06 | 4.4 | L Inferior Frontal Gyrus BA9 |
| | | -40 | 6 | 30 | 2E-02 | 7E-05 | 3.8 | L Precentral Gyrus BA6 |
| | | -42 | 24 | 20 | 2E-02 | 4E-04 | 3.4 | L Middle Frontal Gyrus BA46 |
| 5 | 2456 | 54 | 0 | 46 | 4E-02 | 1E-09 | 6.0 | R Precentral Gyrus BA4 |
| | | 54 | 6 | 36 | 3E-02 | 3E-06 | 4.5 | R Precentral Gyrus BA6 |
| 6 | 1920 | -50 | -6 | 46 | 4E-02 | 3E-10 | 6.2 | L Precentral Gyrus BA4 |
| | | -46 | 0 | 54 | 2E-02 | 2E-04 | 3.6 | L Precentral Gyrus BA6 |
| 7 | 1856 | 0 | 4 | 62 | 3E-02 | 1E-07 | 5.1 | L Medial Frontal Gyrus BA6 |
| *3. PostCG-R BA2: 437 foci, 22 experiments, 310 subjects (x=54, y=-22, z=44)* | | | | | | | | |
| 1 | 5416 | -2 | 4 | 50 | 3E-02 | 2E-09 | 5.9 | L Medial Frontal Gyrus BA6 |
| | | -8 | 12 | 36 | 2E-02 | 1E-05 | 4.2 | L Cingulate Gyrus BA24 |
| 2 | 5312 | 56 | -22 | 44 | 1E-01 | 1E-45 | 14.2 | R Postcentral Gyrus BA2 |
| 3 | 4744 | -42 | -28 | 54 | 3E-02 | 8E-10 | 6.0 | L Inferior Parietal Lobule BA40 |
| | | -34 | -18 | 64 | 3E-02 | 4E-09 | 5.8 | L Precentral Gyrus BA4 |
| 4 | 1528 | 56 | 12 | 36 | 3E-02 | 1E-08 | 5.6 | R Middle Frontal Gyrus BA9 |

| | | | | | | | | |
|---|---|---|---|---|---|---|---|---|
| | | 50 | 8 | 26 | 1E-02 | 4E-04 | 3.3 | R Inferior Frontal Gyrus BA9 |
| 5 | 1432 | 10 | -14 | 6 | 3E-02 | 2E-08 | 5.5 | R Thalamus |
| 6 | 1376 | 56 | 14 | -6 | 3E-02 | 1E-07 | 5.2 | R Superior Temporal Gyrus BA22 |
| 7 | 1376 | 64 | -22 | 18 | 2E-02 | 5E-06 | 4.4 | R Postcentral Gyrus BA40 |
| | | 60 | -16 | 24 | 1E-02 | 3E-04 | 3.4 | R Postcentral Gyrus BA3 |
| 8 | 1056 | 22 | -54 | -22 | 3E-02 | 4E-09 | 5.8 | R Cerebellum. Culmen |
| *NM>M (GM)* | | | | | | | | |
| *4. PreCG-R BA4: 233 foci, 15 experiments, 197 subjects (x=64, y=-14, z=8)* | | | | | | | | |
| 1 | 4000 | 64 | -16 | 38 | 7E-02 | 4E-30 | 11.4 | R Postcentral Gyrus BA3 |
| 2 | 1152 | 34 | 20 | -2 | 2E-02 | 2E-06 | 4.7 | R Claustrum |
| | | 30 | 24 | -12 | 1E-02 | 3E-05 | 4.0 | R Insula BA47 |
| *M>NM (WM)* | | | | | | | | |
| *5. IC/THA-R: 598 foci, 22 experiments, 286 subjects (x=22, y=-14, z=6), nearest grey matter: Right Thalamus.* | | | | | | | | |
| 1 | 8128 | 22 | -14 | 8 | 9E-02 | 2E-40 | 13.3 | R Thalamus |
| | | 38 | 6 | 4 | 2E-02 | 4E-05 | 3.9 | R Claustrum |
| | | 20 | 6 | 6 | 2E-02 | 4E-05 | 3.9 | R Lentiform Nucleus |
| 2 | 6848 | 4 | -2 | 66 | 3E-02 | 8E-09 | 5.6 | R Medial Frontal Gyrus BA6 |
| | | 2 | 8 | 50 | 3E-02 | 3E-07 | 5.0 | L Medial Frontal Gyrus BA6 |
| | | -8 | -6 | 62 | 3E-02 | 5E-07 | 4.9 | L Medial Frontal Gyrus BA6 |
| | | 2 | 12 | 44 | 2E-02 | 2E-05 | 4.1 | L Medial Frontal Gyrus BA32 |
| | | -2 | 2 | 38 | 2E-02 | 3E-05 | 4.1 | L Cingulate Gyrus BA24 |
| | | -2 | 12 | 40 | 2E-02 | 3E-05 | 4.0 | L Cingulate Gyrus BA32 |
| | | 4 | 20 | 32 | 2E-02 | 7E-05 | 3.8 | R Cingulate Gyrus BA32 |
| 3 | 5056 | -24 | -10 | 0 | 3E-02 | 1E-07 | 5.2 | L Lentiform Nucleus |
| | | -20 | -10 | 2 | 3E-02 | 2E-07 | 5.1 | L Lentiform Nucleus |
| | | -12 | 0 | 6 | 2E-02 | 3E-06 | 4.6 | L Thalamus |
| | | -12 | -22 | 4 | 2E-02 | 7E-06 | 4.4 | L Thalamus |
| | | -24 | 0 | -6 | 2E-02 | 6E-05 | 3.9 | L Lentiform Nucleus |
| 4 | 1408 | -28 | -60 | -18 | 3E-02 | 2E-07 | 5.0 | L Cerebellum |
| | | -32 | -60 | -16 | 3E-02 | 3E-07 | 5.0 | L Cerebellum |
| 5 | 1208 | 36 | 22 | 2 | 3E-02 | 1E-07 | 5.2 | R Insula BA13 |
| 6 | 1144 | -8 | -60 | -16 | 3E-02 | 2E-08 | 5.5 | L Cerebellum |
| *NM>M (WM): NA* | | | | | | | | |
| - | - | - | - | - | - | - | - | - |
| **b. FUNCTIONAL ALE ROIs** | | | | | | | | |
| *M>NM* | | | | | | | | |
| *1. IFG-L BA9: 1793 foci, 83 experiments, 1238 subjects (x=-50, y=8, z=18)* | | | | | | | | |
| 1 | 42352 | -50 | 8 | 18 | 3E-01 | 0E+00 | 27.3 | L Inferior Frontal Gyrus BA9 |
| | | -34 | 22 | 0 | 8E-02 | 7E-18 | 8.5 | L Insula BA13 |
| | | -46 | 0 | 46 | 6E-02 | 4E-13 | 7.2 | L Precentral Gyrus BA6 |
| | | -52 | 8 | 36 | 6E-02 | 8E-13 | 7.1 | L Precentral Gyrus BA6 |
| | | -46 | 26 | 16 | 5E-02 | 2E-10 | 6.3 | L Middle Frontal Gyrus BA46 |
| | | -46 | 18 | -6 | 5E-02 | 6E-10 | 6.1 | L Inferior Frontal Gyrus BA47 |
| | | -52 | 30 | -6 | 4E-02 | 2E-08 | 5.5 | L Inferior Frontal Gyrus BA45 |
| | | -42 | 38 | 10 | 4E-02 | 1E-07 | 5.2 | L Middle Frontal Gyrus BA46 |
| | | -18 | 8 | 2 | 4E-02 | 7E-07 | 4.8 | L Lentiform Nucleus |
| | | -32 | -4 | 54 | 4E-02 | 2E-06 | 4.6 | L Precentral Gyrus BA6 |
| | | -36 | -6 | 56 | 3E-02 | 3E-06 | 4.5 | L Precentral Gyrus BA6 |
| | | -18 | 10 | 8 | 3E-02 | 3E-06 | 4.5 | L Lentiform Nucleus |
| | | -24 | 8 | 54 | 3E-02 | 2E-05 | 4.1 | L Sub-Gyral BA6 |
| 2 | 14776 | 50 | 10 | 24 | 8E-02 | 4E-20 | 9.1 | R Inferior Frontal Gyrus BA9 |
| | | 34 | 24 | -2 | 6E-02 | 8E-14 | 7.4 | R Insula BA13 |
| | | 44 | 2 | 44 | 3E-02 | 2E-05 | 4.1 | R Middle Frontal Gyrus BA6 |
| | | 44 | 32 | 20 | 3E-02 | 3E-05 | 4.0 | R Middle Frontal Gyrus BA46 |
| | | 44 | 30 | 24 | 3E-02 | 4E-05 | 3.9 | R Middle Frontal Gyrus BA9 |
| | | 52 | 28 | 24 | 3E-02 | 7E-05 | 3.8 | R Middle Frontal Gyrus BA46 |
| | | 50 | 18 | -6 | 3E-02 | 7E-05 | 3.8 | R Inferior Frontal Gyrus |
| | | 50 | 20 | 10 | 2E-02 | 6E-04 | 3.3 | R Inferior Frontal Gyrus BA44 |
| 3 | 12568 | -30 | -52 | 42 | 7E-02 | 7E-15 | 7.7 | No Gray Matter found |
| | | -48 | -36 | 42 | 6E-02 | 9E-14 | 7.4 | L Inferior Parietal Lobule BA40 |
| | | -24 | -64 | 50 | 5E-02 | 1E-09 | 6.0 | L Superior Parietal Lobule BA7 |
| | | -42 | -44 | 48 | 4E-02 | 1E-08 | 5.6 | L Inferior Parietal Lobule BA40 |
| 4 | 12432 | -4 | 16 | 44 | 6E-02 | 2E-12 | 7.0 | L Medial Frontal Gyrus BA32 |
| | | 6 | 28 | 34 | 5E-02 | 6E-10 | 6.1 | R Cingulate Gyrus BA32 |
| | | -2 | 2 | 56 | 4E-02 | 3E-08 | 5.4 | L Medial Frontal Gyrus BA6 |

| | | | | | | | | |
|---|---|---|---|---|---|---|---|---|
| | | 10 | 16 | 58 | 3E-02 | 1E-04 | 3.6 | R Superior Frontal Gyrus BA6 |
| 5 | 3424 | -46 | -58 | -10 | 5E-02 | 2E-11 | 6.6 | L Fusiform Gyrus BA37 |
| 6 | 3016 | 46 | -38 | 46 | 5E-02 | 1E-09 | 5.9 | R Inferior Parietal Lobule BA40 |
| | | 38 | -56 | 48 | 3E-02 | 4E-05 | 4.0 | R Inferior Parietal Lobule BA7 |
| | | 30 | -60 | 48 | 3E-02 | 4E-05 | 3.9 | R Superior Parietal Lobule BA7 |
| 7 | 2576 | 18 | 10 | 4 | 4E-02 | 4E-08 | 5.4 | R Caudate |
| | | 16 | 6 | 12 | 3E-02 | 1E-05 | 4.3 | R Caudate |
| *2. STG-R BA22: 1793 foci, 83 experiments, 1238 subjects (x=54, y=-10, z=4)* | | | | | | | | |
| 1 | 42352 | -50 | 8 | 18 | 3E-01 | 0E+00 | 27.3 | L Inferior Frontal Gyrus BA9 |
| | | -34 | 22 | 0 | 8E-02 | 7E-18 | 8.5 | L Insula BA13 |
| | | -46 | 0 | 46 | 6E-02 | 4E-13 | 7.2 | L Precentral Gyrus BA6 |
| | | -52 | 8 | 36 | 6E-02 | 8E-13 | 7.1 | L Precentral Gyrus BA6 |
| | | -46 | 26 | 16 | 5E-02 | 2E-10 | 6.3 | L Middle Frontal Gyrus BA46 |
| | | -46 | 18 | -6 | 5E-02 | 6E-10 | 6.1 | L Inferior Frontal Gyrus BA47 |
| | | -52 | 30 | -6 | 4E-02 | 2E-08 | 5.5 | L Inferior Frontal Gyrus BA45 |
| | | -42 | 38 | 10 | 4E-02 | 1E-07 | 5.2 | L Middle Frontal Gyrus BA46 |
| | | -18 | 8 | 2 | 4E-02 | 7E-07 | 4.8 | L Lentiform Nucleus |
| | | -32 | -4 | 54 | 4E-02 | 2E-06 | 4.6 | L Precentral Gyrus BA6 |
| | | -36 | -6 | 56 | 3E-02 | 3E-06 | 4.5 | L Precentral Gyrus BA6 |
| | | -18 | 10 | 8 | 3E-02 | 3E-06 | 4.5 | L Lentiform Nucleus |
| | | -24 | 8 | 54 | 3E-02 | 2E-05 | 4.1 | L Sub-Gyral BA6 |
| 2 | 14776 | 50 | 10 | 24 | 8E-02 | 4E-20 | 9.1 | R Inferior Frontal Gyrus BA9 |
| | | 34 | 24 | -2 | 6E-02 | 8E-14 | 7.4 | R Insula BA13 |
| | | 44 | 2 | 44 | 3E-02 | 2E-05 | 4.1 | R Middle Frontal Gyrus BA6 |
| | | 44 | 32 | 20 | 3E-02 | 3E-05 | 4.0 | R Middle Frontal Gyrus BA46 |
| | | 44 | 30 | 24 | 3E-02 | 4E-05 | 3.9 | R Middle Frontal Gyrus BA9 |
| | | 52 | 28 | 24 | 3E-02 | 7E-05 | 3.8 | R Middle Frontal Gyrus BA46 |
| | | 50 | 18 | -6 | 3E-02 | 7E-05 | 3.8 | R Inferior Frontal Gyrus |
| | | 50 | 20 | 10 | 2E-02 | 6E-04 | 3.3 | R Inferior Frontal Gyrus BA44 |
| 3 | 12568 | -30 | -52 | 42 | 7E-02 | 7E-15 | 7.7 | No Gray Matter found |
| | | -48 | -36 | 42 | 6E-02 | 9E-14 | 7.4 | L Inferior Parietal Lobule BA40 |
| | | -24 | -64 | 50 | 5E-02 | 1E-09 | 6.0 | L Superior Parietal Lobule BA7 |
| | | -42 | -44 | 48 | 4E-02 | 1E-08 | 5.6 | L Inferior Parietal Lobule BA40 |
| 4 | 12432 | -4 | 16 | 44 | 6E-02 | 2E-12 | 7.0 | L Medial Frontal Gyrus BA32 |
| | | 6 | 28 | 34 | 5E-02 | 6E-10 | 6.1 | R Cingulate Gyrus BA32 |
| | | -2 | 2 | 56 | 4E-02 | 3E-08 | 5.4 | L Medial Frontal Gyrus BA6 |
| | | 10 | 16 | 58 | 3E-02 | 1E-04 | 3.6 | R Superior Frontal Gyrus BA6 |
| 5 | 3424 | -46 | -58 | -10 | 5E-02 | 2E-11 | 6.6 | L Fusiform Gyrus BA37 |
| 6 | 3016 | 46 | -38 | 46 | 5E-02 | 1E-09 | 5.9 | R Inferior Parietal Lobule BA40 |
| | | 38 | -56 | 48 | 3E-02 | 4E-05 | 4.0 | R Inferior Parietal Lobule BA7 |
| | | 30 | -60 | 48 | 3E-02 | 4E-05 | 3.9 | R Superior Parietal Lobule BA7 |
| 7 | 2576 | 18 | 10 | 4 | 4E-02 | 4E-08 | 5.4 | R Caudate |
| | | 16 | 6 | 12 | 3E-02 | 1E-05 | 4.3 | R Caudate |
| *3. STG-L BA22: 588 foci, 36 experiments, 555 subjects (x=-58, y=-46, z=16)* | | | | | | | | |
| 1 | 8320 | -58 | -44 | 16 | 1E-01 | 0E+00 | 16.8 | L Superior Temporal Gyrus BA22 |
| 2 | 4272 | -44 | 24 | 20 | 3E-02 | 6E-07 | 4.9 | L Middle Frontal Gyrus BA46 |
| | | -44 | 14 | 22 | 2E-02 | 2E-06 | 4.6 | L Inferior Frontal Gyrus BA9 |
| | | -42 | 12 | 28 | 2E-02 | 6E-06 | 4.4 | L Inferior Frontal Gyrus BA9 |
| | | -40 | 2 | 44 | 2E-02 | 2E-05 | 4.1 | L Precentral Gyrus BA6 |
| | | -44 | 6 | 20 | 2E-02 | 3E-05 | 4.1 | L Inferior Frontal Gyrus BA9 |
| | | -40 | 0 | 32 | 2E-02 | 3E-05 | 4.0 | L Precentral Gyrus BA6 |
| 3 | 2368 | 58 | -38 | 4 | 3E-02 | 2E-07 | 5.0 | R Middle Temporal Gyrus BA22 |
| | | 50 | -38 | 6 | 2E-02 | 1E-05 | 4.2 | R Superior Temporal Gyrus BA41 |
| | | 56 | -50 | 6 | 2E-02 | 1E-05 | 4.2 | R Middle Temporal Gyrus BA21 |
| | | 62 | -42 | 16 | 2E-02 | 1E-04 | 3.7 | R Superior Temporal Gyrus BA13 |
| | | 48 | -32 | 10 | 2E-02 | 3E-04 | 3.4 | R Superior Temporal Gyrus BA41 |
| 4 | 2272 | 34 | 20 | 0 | 3E-02 | 7E-07 | 4.8 | R Claustrum |
| | | 48 | 16 | 4 | 2E-02 | 3E-05 | 4.0 | R Precentral Gyrus BA44 |
| | | 52 | 26 | -2 | 2E-02 | 2E-04 | 3.5 | R Inferior Frontal Gyrus BA45 |
| 5 | 2272 | -42 | 16 | 2 | 3E-02 | 2E-09 | 5.9 | L Insula BA13 |
| | | -52 | 24 | 6 | 2E-02 | 1E-05 | 4.2 | L Inferior Frontal Gyrus BA45 |
| 6 | 1680 | -6 | 12 | 50 | 3E-02 | 2E-07 | 5.1 | L Medial Frontal Gyrus BA6 |
| *NM>M* | | | | | | | | |
| *4. IPL-L BA40: 565 foci, 28 experiments, 321 subjects (x=-50, y=-36, z=56)* | | | | | | | | |
| 1 | 10416 | -48 | -36 | 54 | 1E-01 | 0E+00 | 14.6 | L Inferior Parietal Lobule BA40 |

| | | | | | | | | |
|---|---|---|---|---|---|---|---|---|
| | | -40 | -12 | 56 | 2E-02 | 1E-07 | 5.2 | L Precentral Gyrus BA4 |
| | | -34 | -28 | 64 | 2E-02 | 8E-05 | 3.8 | L Postcentral Gyrus BA3 |
| 2 | 4808 | -2 | 6 | 54 | 3E-02 | 2E-09 | 5.9 | L Medial Frontal Gyrus BA6 |
| | | -4 | -6 | 68 | 1E-02 | 7E-04 | 3.2 | L Medial Frontal Gyrus BA6 |
| 3 | 1720 | 52 | 20 | -4 | 2E-02 | 1E-07 | 5.2 | R Inferior Frontal Gyrus BA47 |
| 4 | 1376 | -58 | 4 | 30 | 2E-02 | 3E-06 | 4.5 | L Precentral Gyrus BA6 |
| | | -50 | 10 | 30 | 2E-02 | 5E-05 | 3.9 | L Inferior Frontal Gyrus BA9 |
| *5. PreCG-L BA6: 2068 foci, 108 experiments, 1982 subjects (x=-48, y=8, z=36)* | | | | | | | | |
| 1 | 33856 | -48 | 8 | 36 | 4E-01 | 0E+00 | 29.9 | L Precentral Gyrus BA6 |
| | | -32 | 22 | 0 | 1E-01 | 6E-28 | 10.9 | L Claustrum |
| | | -48 | 24 | 26 | 6E-02 | 2E-12 | 6.9 | L Middle Frontal Gyrus BA9 |
| | | -52 | 10 | 16 | 5E-02 | 4E-09 | 5.8 | L Inferior Frontal Gyrus BA44 |
| | | -50 | 12 | 2 | 5E-02 | 6E-09 | 5.7 | L Insula BA13 |
| | | -26 | -4 | 58 | 5E-02 | 4E-08 | 5.4 | L Middle Frontal Gyrus BA6 |
| | | -50 | 28 | 4 | 4E-02 | 2E-07 | 5.1 | L Inferior Frontal Gyrus BA45 |
| | | -46 | 24 | -6 | 4E-02 | 9E-07 | 4.8 | L Inferior Frontal Gyrus BA47 |
| | | -50 | 14 | -14 | 4E-02 | 8E-06 | 4.3 | L Superior Temporal Gyrus BA38 |
| 2 | 15696 | -24 | -66 | 46 | 7E-02 | 6E-14 | 7.4 | L Precuneus BA7 |
| | | -36 | -52 | 46 | 7E-02 | 1E-13 | 7.3 | L Inferior Parietal Lobule BA40 |
| | | -44 | -36 | 48 | 4E-02 | 6E-07 | 4.9 | L Inferior Parietal Lobule BA40 |
| 3 | 14160 | -4 | 14 | 50 | 1E-01 | 9E-27 | 10.6 | L Superior Frontal Gyrus BA6 |
| | | 6 | 28 | 36 | 6E-02 | 2E-10 | 6.3 | R Cingulate Gyrus BA32 |
| | | -6 | 24 | 32 | 4E-02 | 4E-06 | 4.5 | L Cingulate Gyrus BA32 |
| | | -4 | 32 | 28 | 3E-02 | 2E-05 | 4.2 | L Cingulate Gyrus BA32 |
| 4 | 11032 | 50 | 10 | 26 | 8E-02 | 1E-16 | 8.2 | R Inferior Frontal Gyrus BA9 |
| | | 48 | 36 | 24 | 4E-02 | 2E-07 | 5.1 | R Middle Frontal Gyrus BA9 |
| 5 | 5952 | 40 | -50 | 48 | 5E-02 | 9E-10 | 6.0 | R Inferior Parietal Lobule BA40 |
| | | 34 | -60 | 46 | 5E-02 | 2E-09 | 5.9 | R Precuneus BA19 |
| | | 46 | -38 | 50 | 4E-02 | 6E-06 | 4.4 | R Inferior Parietal Lobule BA40 |
| | | 26 | -68 | 48 | 3E-02 | 2E-05 | 4.1 | R Precuneus BA7 |
| | | 24 | -70 | 52 | 3E-02 | 2E-05 | 4.1 | R Precuneus BA7 |
| | | 26 | -58 | 60 | 3E-02 | 6E-04 | 3.2 | R Superior Parietal Lobule BA7 |
| 6 | 5656 | 34 | 22 | -2 | 9E-02 | 6E-20 | 9.1 | R Insula |
| | | 52 | 18 | -6 | 3E-02 | 3E-05 | 4.0 | R Inferior Frontal Gyrus |
| 7 | 4464 | -44 | -60 | -16 | 5E-02 | 1E-09 | 6.0 | L Fusiform Gyrus BA37 |
| | | -40 | -72 | -12 | 5E-02 | 1E-08 | 5.6 | L Fusiform Gyrus BA19 |
| | | -50 | -54 | 0 | 3E-02 | 2E-04 | 3.5 | L Middle Temporal Gyrus BA37 |
| 8 | 3864 | -8 | -12 | 8 | 5E-02 | 3E-09 | 5.8 | L Thalamus |
| | | -18 | 8 | 4 | 3E-02 | 4E-05 | 3.9 | L Lentiform Nucleus |
| 9 | 1800 | 32 | 0 | 50 | 4E-02 | 4E-07 | 5.0 | R Middle Frontal Gyrus BA6 |

ALE, anatomic likelihood estimation; M, musicians; NM, non-musicians; GM, gray matter; WM, white matter; BA, Brodmann area; ROIs, regions-of-interest; P, p-value; Z, peak z-value; R, right; L, left. **ROIs**: IFG, inferior frontal gyrus; IPL, inferior parietal lobule; IC, internal capsule; PostCG, postcentral gyrus (primary somatosensory cortex or S1); PreCG, precentral gyrus (primary motor cortex or M1); STG, superior temporal gyrus (primary auditory cortex). Music-related ROIs were created in Mango (http://rii.uthscsa.edu/mango//userguide.html) with a 5mm-radius sphere. Last search in Sleuth, 10.10.2021 (http://www.brainmap.org/sleuth/); NA, not enough available observations.

**Supplementary Table 5. Functional characterization of brain regions resulted from structural and functional ALE meta-analyses.**

| | |
|---|---|
| **a. STRUCTURAL ALE ROIs** | |
| *M>NM (GM)* | |
| *1. STG-R BA13: 992 foci, 46 experiments, 593 subjects (x=50, y=-20, z=8)* | |
| Action | Execution, speech, motor learning, preparation |
| Cognition | Attention, language, semantics speech, syntax, explicit memory, working memory, music, reasoning |
| Emotion | Anger, anxiety, sadness, positive emotion, happiness, reward/gain |
| Interoception | Sexuality |
| Perception | Audition, somesthesis, pain, vision |
| Paradigms | Affective words, classical conditioning, counting/calculation, delayed match to sample, emotion induction, face discrimination, finger tapping/button pressing, flexion/extension, go/no-go, meditation, music comprehension, music production, n-back, naming overt, pain discrimination, passive listening, passive viewing, phonological discrimination, pitch discrimination, reading overt, reasoning/problem solving, recitation/repetition, reward, semantic discrimination, sequence recall/learning, sexual arousal, syntactic discrimination, tone discrimination, word generation |
| *2. STG-L BA41: 1428 foci, 71 experiments, 961 subjects (x=-56, y=-20, z=2)* | |
| Action | Execution, speech, imagination, inhibition, motor learning, observation, preparation |
| Cognition | Attention, language, orthography, phonology, semantics, speech, syntax, explicit memory, working memory, music, reasoning, spatial |
| Emotion | Disgust, fear, guilt, sadness, happiness, valence |
| Interoception | Sexuality, thermoregulation |
| Perception | Audition, pain, vision, motion, shape |
| Paradigms | Cued explicit recognition, emotion induction, face discrimination, film viewing, finger tapping/button pressing, free list word recall, go/no-go, imagined objects/scenes, mental rotation, music comprehension, music production, naming, oddball discrimination, orthographic discrimination, pain discrimination, paired associate recall, passive listening, passive viewing, phonological discrimination, pitch discrimination, reading, reasoning/problem solving, recitation/repetition, semantic discrimination, sequence recall/learning, sexual arousal, tone discrimination, visual motion, visuospatial attention, word generation |
| *3. PostCG-R BA2: 437 foci, 22 experiments, 310 subjects (x=54, y=-22, z=44)* | |
| Action | Execution, motor learning |
| Cognition | Attention, somatic |
| Emotion | - |
| Interoception | Respiration regulation |
| Perception | Audition, somesthesis, pain, vision, colour, shape |
| Paradigms | Chewing/swallowing, drawing, face discrimination, finger tapping/button pressing, flanker, flexion/extension, go/no-go, hypercapnia, motor learning, oddball discrimination, pain discrimination, passive listening, passive viewing, pursuit/tracking, saccades, tactile discrimination, tone discrimination, transcranial magnetic stimulation, visuospatial attention, writing |
| *NM>M (GM)* | |
| *4. PreCG-R BA4: 233 foci, 15 experiments, 197 subjects (x=64, y=-14, z=8)* | |
| Action | Execution, inhibition, observation, preparation |
| Cognition | Attention, semantics, speech, social cognition, temporal |
| Emotion | Sadness, happiness |
| Interoception | - |
| Perception | Gustation, somesthesis, pain, vision |
| Paradigms | Competition/cooperation, deception, face discrimination, film viewing, finger tapping/button pressing, flanker, flexion/extension, go/no-go, grasping, imagined movement, isometric force, pain discrimination, passive viewing, recitation/repetition, taste, transcranial magnetic stimulation, video games, visuospatial attention |
| *M>NM (WM)* | |
| *5. IC-R: 598 foci, 22 experiments, 286 subjects (x=22, y=-14, 6), nearest grey matter: Right Thalamus.* | |
| Action | Execution, speech, imagination |
| Cognition | Attention, orthography, explicit memory, working memory, reasoning |
| Emotion | Negative emotion, happiness, reward |
| Interoception | Thermoregulation |
| Perception | Audition, olfaction, pain |
| Paradigms | Counting/calculation, cued explicit recognition/recall, emotion induction, episodic recall, face discrimination, finger tapping/button pressing, flexion/extension, free list word recall, go/no-go, imagined movement, isometric force, n-back, olfactory discrimination, pain discrimination, passive listening, reading, recitation/repetition, reward, tone discrimination |
| *NM>M (WM): NA* | |
| - | |

| **b. FUNCTIONAL ALE ROIs** | |
|---|---|
| *M>NM* | |
| *1. IFG-L BA9: 1793 foci, 83 experiments, 1238 subjects (x=-50, y=8, z=18)* | |
| Action | Execution, speech, imagination, inhibition, observation, preparation |
| Cognition | Attention, language, orthography, phonology, semantics, speech, syntax, explicit memory, working memory, music, reasoning, social cognition, spatial |
| Emotion | Negative emotion, anger, fear, reward |
| Interoception | - |
| Perception | Audition, gustation, somesthesis, pain, vision, motion, shape |
| Paradigms | Chewing/swallowing, counting/calculation, cued explicit recognition/recall, delayed match to sample, driving, encoding, face discrimination, figurative language, film viewing, finger tapping/button pressing, flexion/extension, gambling, go/no-go, imagined movement, imagined objects/scenes, lexical decision, magnitude comparison, meditation, mental rotation, music comprehension, music production, n-back, naming, orthographic discrimination, pain discrimination, paired associate recall, passive listening, passive viewing, phonological discrimination, reading, reasoning/problem solving, recitation/repetition, reward, saccades, semantic discrimination, sequence recall/learning, tactile discrimination, theory-of-mind, tone discrimination, visual object identification, Wisconsin Card Sorting Test, word generation, word stem completion |
| *2. STG-R BA22: 1793 foci, 83 experiments, 1238 subjects (x=54, y=-10, z=4)* | |
| Action | Speech, imagination, inhibition, motor learning |
| Cognition | Attention, phonology, speech, music, reasoning, spatial |
| Emotion | Reward |
| Interoception | Hunger, osmoregulation, thermoregulation, thirst |
| Perception | Audition, gustation, somesthesis, pain, vision |
| Paradigms | Acupuncture, counting/calculation, emotion induction, encoding, face discrimination, film viewing, finger tapping/button pressing, flexion/extension, gambling, go/no-go, hunger, imagined movement, imagined objects/scenes, multi-tasking, music comprehension, music production, pain discrimination, passive listening, passive viewing, phonological discrimination, pitch discrimination, reading, reasoning/problem solving, reward, semantic discrimination, sequence recall/learning, taste, thirst induction, tone discrimination |
| *3. STG-L BA22: 588 foci, 36 experiments, 555 subjects (x=-58, y=-46, z=16)* | |
| Action | Execution, speech, imagination, inhibition, observation |
| Cognition | Attention, language, phonology, semantics, speech, explicit memory, music, reasoning, social cognition |
| Emotion | Disgust, embarrassment, positive emotion |
| Interoception | Sexuality |
| Perception | Audition, somesthesis, pain, vision |
| Paradigms | Acupuncture, affective pictures, controlled breathing, cued explicit recognition/recall, divided auditory attention, emotion induction, emotional body language perception, encoding, face discrimination, finger tapping/button pressing, go/no-go, music comprehension, music production, oddball discrimination, pain discrimination, paired associate recall, passive viewing, phonological discrimination, pitch discrimination, reading, reasoning/problem solving, recitation/repetition, semantic discrimination, Stroop-color, theory-of-mind, tone discrimination, visual motion, visuospatial attention, word generation |
| *NM>M* | |
| *4. IPL-L BA40: 565 foci, 28 experiments, 321 subjects (x=-50, y=-36, z=56)* | |
| Action | Execution, speech, imagination, inhibition, motor learning, preparation |
| Cognition | Attention, orthography, phonology, semantics, explicit memory, working memory, music, reasoning, social cognition, spatial |
| Emotion | Negative emotion, anxiety, reward |
| Interoception | - |
| Perception | Audition, somesthesis, pain, vision, shape |
| Paradigms | Classical conditioning, counting/calculation, cued explicit recognition/recall, emotion induction, film viewing, finger tapping/button pressing, fixation, flexion/extension, go/no-go, imagined movement, imagined objects/scenes, mental rotation, motor learning, multi-tasking, object discrimination, pain discrimination, phonological discrimination, reading, reasoning/problem solving, recitation/repetition, reward, semantic discrimination, Stroop colour, tactile discrimination, task switching, tone discrimination, visual object identification, visuospatial attention |
| *5. PreCG-L BA6: 2068 foci, 108 experiments, 1982 subjects (x=-48, y=8, z=36)* | |
| Action | Execution, speech, imagination, inhibition, motor learning, observation, preparation |
| Cognition | Attention, language, orthography, phonology, semantics, speech, syntax, explicit memory, working memory, music, reasoning, social cognition, somatic, spatial, temporal |
| Emotion | Negative emotion, anger, disgust, far, positive emotion, happiness, reward |
| Interoception | Hunger, sexuality, sleep |
| Perception | Audition, gustation, somesthesis, vision, color, motion, shape |
| Paradigms | Affective pictures, anti-saccades, chewing/swallowing, counting/calculation, cued explicit recognition/recall, deception, delayed match to sample, divided auditory attention, drawing, driving, emotion induction, encoding, episodic recall, estimation, face discrimination, figurative language, film viewing, finger tapping/button pressing, flanker, gambling, go/no-go, hunger, imagined movement, imagined objects/scenes, mental rotation, multi-tasking, music comprehension, |

| | |
|---|---|
| | n-back, naming, oddball discrimination, orthographic discrimination, paired associate recall, phonological discrimination, pitch discrimination, pursuit, reading, reasoning/problem solving, recitation, reward, saccades, semantic discrimination, Stroop, tactile discrimination, task switching, taste, theory of mind, tone discrimination, visual object identification, visual pursuit, visuospatial attention, Wisconsin card sorting, word generation |

ALE, anatomic likelihood estimation; M, musicians; NM, non-musicians; GM, grey matter; WM, white matter; BA, Brodmann area; ROIs, regions-of-interest; P, p-value; Z, peak z-value; R, right; L, left. **ROIs**: IFG, inferior frontal gyrus; IPL, inferior parietal lobule; IC, internal capsule; PostCG, postcentral gyrus (primary somatosensory cortex or S1); PreCG, precentral gyrus (primary motor cortex or M1); STG, superior temporal gyrus (primary auditory cortex). Music-related ROIs were created in Mango (http://rii.uthscsa.edu/mango//userguide.html) with a 5mm-radius sphere. Last search in Sleuth, 10.10.2021 (http://www.brainmap.org/sleuth/); NA, not enough available observations.

**Supplementary Table 6. FSN robustness assessment of brain regions resulted from structural and functional ALE meta-analyses.**

| Cluster number | Volume ($mm^3$) | MNI coordinates x | y | z | ALE | Label (Side, region) | Contributing studies (k) | FSN |
|---|---|---|---|---|---|---|---|---|
| **a. STRUCTURAL ALE ROIs** | | | | | | | | |
| *M>NM (GM): 133 foci, 20 experiments, 1071 subjects,* minimum FSN = 6 | | | | | | | | |
| 1 | 912 | 50 | -20 | 8 | 2E-02 | R Superior Temporal Gyrus BA13 | 5 | 10 |
| 2 | 784 | -56 | -20 | 2 | 2E-02 | L Superior Temporal Gyrus BA41 | 3 | 10 |
| 3 | 536 | 54 | -22 | 44 | 2E-02 | R Postcentral Gyrus BA2 (S1) | 3 | <6 |
| *NM>M (GM): 22 foci, 8 experiments, 305 subjects,* minimum FSN = 3 | | | | | | | | |
| 4 | 520 | 64 | -14 | 38 | 1E-02 | R Precentral Gyrus BA4 (MI) | 2 | <3 |
| *M>NM (WM): 22 foci, 5 experiments, 139 subjects,* minimum FSN = 2 | | | | | | | | |
| 5 | 792 | 22 | -14 | 6 | 1E-02 | R Thalamus, Internal Capsule | 3 | 18 |
| *NM>M (WM): NA* | | | | | | | | |
| - | - | - | - | - | - | - | - | - |
| **b. FUNCTIONAL ALE ROIs** | | | | | | | | |
| *M>NM: 354 foci, 34 experiments, 979 subjects,* minimum FSN = 10 | | | | | | | | |
| 1 | 3080 | -50 | 8 | 18 | 3E-02 | L Inferior Frontal Gyrus BA9 | 24 | 90 |
| 2 | 1136 | 54 | -10 | 4 | 2E-02 | R Superior Temporal Gyrus BA22 | 7 | 70 |
| 3 | 920 | -58 | -46 | 16 | 2E-02 | L Superior Temporal Gyrus BA22 | 7 | <10 |
| *NM>M: 144 foci, 12 experiments, 286 subjects,* minimum FSN = 4 | | | | | | | | |
| 4 | 912 | -50 | -36 | 56 | 1E-02 | L Inferior Parietal Lobule BA40 | 5 | 8 |
| 5 | 736 | -48 | 8 | 36 | 2E-02 | L Precentral Gyrus BA6 | 5 | <4 |

FSN, Fail-Safe N analysis; NA, not enough available observations.

[Citations of included studies](#)

**Abbreviations**

| | |
|---|---|
| ACC | anterior cingulate cortex |
| AF | arcuate fasciculus |
| AnG | angular gyrus |
| CalC | calcarine cortex |
| CC | corpus callosum |
| CAU | caudate |
| CLAU | claustrum |
| CRBL | cerebellum |
| CST | corticospinal tract |
| CUN | cuneus |
| DLPFC | dorsolateral prefrontal cortex |
| EC | entorhinal cortex |
| Fmaj | forceps major |
| Fmin | forceps minor |
| FO | frontal operculum |
| FusG | fusiform gyrus |
| GP | globus pallidus |
| HG | Helsch's gyrus |
| HIPP | hippocampus |
| IC | internal capsule |
| IF | inferior colliculus |
| IFG | inferior frontal gyrus |
| IFOF | inferior fronto-occipital fasciculus |
| ILF | inferior longitudinal fasciculus |
| INS | insula |
| IPL | inferior parieal lobule |
| ITG | inferior temporal gyrus |
| LG | lingual gyrus |
| LOC | lateral occipital cortex |
| MB | midbrain |
| MCC | middle cingulate cortex |
| MCP | middle cerebellar peduncle |
| MedFG | medial frontal gyrus |
| MidFG | middle frontal gyrus |
| MidTG | middle temporal gyrus |
| OFC | orbitofrontal cortex |
| PaHIPP | parahippocampal gyrus |
| PCC | posterior cingulate cortex |
| PCN | precuneus |
| PMC | premotor cortex |
| PO | parietal operculum |
| PostCG | postcentral gyrus (primary somatosensory cortex or SI) |
| PP | planum polare |
| PreCG | precentral gyrus (primiary motor cortex or M1) |
| PT | planum temporale |
| PUT | putamen |
| RN | red nucleus |
| SCP | superior cerebellar peduncle |
| SFG | superior frontal gyrus |
| SII | secondary somatosensory cortex |
| SLF | superior longitudial fasciculus |
| SMA | supplementary motor area |
| SMG | supramarginal gyrus |
| SPL | superior parietal lobule |
| STG | superior temporal gyrus |
| STS | superior temporal sulcus |
| THA | thalamus |
| TP | temporal pole |
| TPG | temporoparietal junction |
| VER | Vermis |
